# Not all sexual monomorphisms evolve equally: female and male ornamentation in Tyranni passerines

**DOI:** 10.1101/2025.09.23.678134

**Authors:** Gabriel Macedo, Yuchen Chen, Sara Lipshutz

**Affiliations:** Duke University

**Keywords:** social selection, sexual selection, mutual mate choice, spectrophotometry, coloration

## Abstract

Female animals often have ornaments such as conspicuous coloration. However, the evolution of female ornamentation and its contribution to sexual dimorphism are understudied relative to male ornamentation. We addressed this knowledge gap by investigating plumage ornamentation of Tyranni (Suboscines). We hypothesized that greater ornamentation is associated with stronger sociosexual selection in females and males, and that sexual dimorphism emerges when selection pressures differ between sexes. We tested associations of ornamentation and sexual dimorphism with territoriality, mating system, parental care, nest type, migratory behavior, and climatic seasonality. We found that females and males show similarly high ornamentation (elaborate monomorphism) in territorial, non-migratory, monogamous species with biparental care, and species nesting in cavities. In contrast, males show higher ornamentation than females in migratory and polygynous species with female-only parental care and in more seasonal climates, though these cases were less common than elaborate monomorphism. Our findings suggest that when the sexes share sociosexual selection pressures, they show similar ornamentation, whether elaborate or inconspicuous. Conversely, sexual dimorphism is greater when males face stronger sociosexual selection pressures than females. Our study provides a framework for testing how sociosexual selection shapes female traits, which is key to understanding the evolution of sex differences and similarities.

## Introduction

Research on the evolution of elaborate traits such as ornamentation has traditionally focused on male-biased sexual dimorphism, in which traits are more exaggerated in males than in females (Ah-King, 2022; Andersson, 1994; Bateman, 1948; Darwin, 1871). This emphasis has been motivated by theoretical and empirical studies suggesting that sexual selection is predominantly driven by male-male competition and female mate choice (Arnold & Duvall, 1994; Bateman, 1948; Darwin, 1871). However, the presence of female ornamentation across diverse taxa poses some challenges for a strictly male-biased sexual selection framework (Amundsen, 2000; Bonduriansky, 2001, 2009; Fromonteil et al., 2023; Hare & Simmons, 2019; Nolazco et al., 2022; Odom et al., 2025). Historic hypotheses proposed that female ornamental traits result from genetic or developmental correlations between the sexes, as well as weak sexual selection pressures (Darwin, 1871; Lande, 1980). However, growing evidence indicates that females across taxa use ornamental traits to compete for mates (Hare & Simmons, 2019; Murray et al., 2020), breeding territories (Cain & Langmore, 2016; Rosvall, 2008), and food (Falk et al., 2021), indicating that female ornaments are shaped by direct selection pressures. Despite the prevalence of female ornaments, their evolutionary drivers are consistently understudied in relation to their male counterparts (Ah-King, 2022; Nolazco et al., 2022).

Along a continuum, the degree of variation between female and male traits can range from extreme sex differences (male- or female-biased sexual dimorphism) to similarities (sexual monomorphism). Sexual monomorphism can be further understood as elaborate (“bright”) monomorphism (when sexes are similarly ornamented) or inconspicuous (“dull”) monomorphism (when sexes are similarly inconspicuous; (Kraaijeveld et al., 2007). Examples of elaborate monomorphism include coloration, songs, and courtship behaviour across diverse taxa (Beco et al., 2021; Egger et al., 2008; Greenquist, 1982; Masello & Quillfeldt, 2003; Odom et al., 2025; Rull et al., 2016; Tigreros et al., 2014), in which females and males show similarly elaborate signals that function in contexts such as mutual mate choice, pair-bond maintenance, and assortative mating. In turn, examples of inconspicuous monomorphism include species in which females and males show similarly cryptic coloration associated with camouflage against predators (Baños-Villalba et al., 2018; Diniz et al., 2016). Despite evolving through different processes, elaborate and inconspicuous monomorphisms are often conflated, that is, studies often assume that all monomorphic taxa are under weak sexual selection pressures (Kraaijeveld et al., 2007).

Social selection theory provides a comprehensive framework to address the evolution of sex differences and similarities (Crook, 1972; West-Eberhard, 1979, 1983, 2014). Social selection (henceforth referred to as sociosexual selection) is defined as the “differential survival and reproduction resulting from differential success in social competition” (West-Eberhard, 1979, 1983, 2014). When in temporary or permanent social groups, individuals, regardless of their sex, may use traits such as ornaments and weapons to compete over mates and fertilizations (which corresponds to sexual selection *sensu strictu*; (Darwin, 1871; West-Eberhard, 1983) and other key resources such as territories, nest sites, and foraging grounds (Lyon & Montgomerie, 2012; West-Eberhard, 1983). Sociosexual selection, therefore, is able to address the evolution of elaborate traits in either sex under sex-biased sexual selection, mutual sexual selection (Kraaijeveld et al., 2007), as well as selection resulting from non-sexual aspects of social competition (Lyon & Montgomerie, 2012; West-Eberhard, 1983).

Predictions concerning sociosexual selection and the evolution of sex differences and similarities were laid out decades ago (West-Eberhard, 1979, 1983) However, empirical studies addressing evolutionary drivers of both female and male traits have been limited (Ah-King, 2022; Hare & Simmons, 2019; Nolazco et al., 2022; Riebel et al., 2019). A specific prediction of sociosexual selection theory is that when one sex competes with greater intensity than the other, sexual dimorphism of social signals should evolve. In contrast, when females and males socially compete with similarly high intensities, elaborate monomorphism should evolve (Kraaijeveld et al., 2007; West-Eberhard, 1983). Similarly, inconspicuous monomorphism should evolve when females and males are mostly solitary and only weakly engage in social competition (Kraaijeveld et al., 2007; West-Eberhard, 1983). Natural selection pressures must also be considered, as selection unrelated to social signaling and competition can also affect traits such as coloration (West-Eberhard, 1979, 1983). For instance, elaborate or inconspicuous monomorphism might evolve under predation pressure through aposematism or camouflage, respectively (Gade et al., 2016; Medina et al., 2017).

Birds have been a key system to test hypotheses about the evolution of elaborate social traits (Darwin, 1871; West-Eberhard, 1983). Birds show a large diversity of social signals that vary widely in their degree of sex differences and elaboration, especially in plumage with varying color saturation, diversity and/or contrast (Barber et al., 2024; Cooney et al., 2022; Dale et al., 2015). In some cases, the sexes are so differently colored that they have even been mistaken as being from different species, as with the brightly colored females and cryptic males of *Eclectus* parrots (Forbes & Forbes, 1877; Heinsohn et al., 2005; LeBas, 2006). In other cases, precise spectrophotometric measurements reveal that the sexes are similarly dull and inconspicuously colored (Diniz et al., 2016; Marcondes & Brumfield, 2019). Moreover, birds occupy diverse habitats and ecological niches and show large variation in life history traits that affect social competition, such as mating systems and territorial behavior (Drury et al., 2024; Tobias & Pigot, 2019). For instance, stronger social competition is generally associated with polygamous mating systems and territoriality (Barber et al., 2024; Dale et al., 2015). Due to the diversity of life history traits and sexual variation in signals such as plumage color, birds constitute an excellent system to address the factors shaping the evolution of female traits and the consequences for sex differences and similarities.

Here, we use the plumage color ornamentation (i.e., color saturation, diversity or contrast) of passerines as a model system of elaborate social traits to ask: (1) Do sociosexual selection pressures such as mating system, territoriality and pair bonding shape plumage ornamentation similarly in females and males? (2) Do natural selection pressures such as vulnerability to predation affect female and male plumage elaboration similarly or differently? Based on (West-Eberhard, 1983), we predicted that female trait elaboration is associated with stronger social competition, and elaborate monomorphism is associated with similar intensities of social competition in females and males. Additionally, we predicted that natural selection pressures affect females more strongly when female investment in parental care is greater than that of males (Andersson, 1994; Trivers, 1972). Addressing the factors shaping female trait elaboration is key to obtaining a comprehensive understanding of the evolution of sex differences and similarities.

## Methods

### Focal taxa

We focused on Tyranni passerines (Suboscines), one of the largest tropical bird radiations (Harvey et al., 2020). Some Tyranni species show elaborately monomorphic plumages such as tyrant flycatchers (Winkler et al., 2020), whereas others are inconspicuously monomorphic in plumage coloration, such as many ovenbirds (Marcondes & Brumfield, 2019). There are classic examples of male-biased sexual dichromatism (i.e., brightly colored males and cryptic females) such as in cotingas and manakins (Berv & Prum, 2014; Cooney et al., 2019), and less known cases of female-biased sexual dichromatism such in antbirds (Zimmer & Isler, 2003). Additionally, mating systems, parental care, and territoriality vary widely in Tyranni. Many socially monogamous species form long-term social pairs that share parental care and defend territories year-round, whereas others are non-territorial, lek-forming, and socially polygynous with female-only parental care (Berv & Prum, 2014; Cooney et al., 2019; Harvey et al., 2020; Remsen, 2003; Zimmer & Isler, 2003), although genetic polyandry has been recorded even in species previously considered strictly polygynous such as the manakins (Gaiotti et al., 2020).

### Plumage color elaboration

We obtained data on the plumage coloration of species of Tyranni from Marcondes & Brumfield (2019), which included the parvorder Furnariida, and Cooney et al., 2019, which included the parvorder Tyrannida. Together, Furnariida and Tyrannida comprise a monophyletic clade (Infraorder Tyrannides) and most species of Tyranni (Harvey et al., 2020). Marcondes & Brumfield (2019) used a spectrophotometer to measure plumage reflectance of museum specimens in six plumage patches (belly, breast, throat, wing, crown, and tail) on an average of 3.2 males and 2.9 females per species. Cooney et al. (2019) measured plumage coloration based on calibrated digital photographs of museum specimens mapped into avian cone-catch values on an average of 2.7 males and 2.4 females per species. Photograph-based color estimates were highly correlated with spectrophotometric data (Pearson’s r > 0.9 across cone types), providing a validated method of estimating plumage coloration across studies (Cooney et al., 2019). We extracted from Cooney et al. 2019 measurements from the same plumage patches as in Marcondes and Brumfield 2019. We removed species that did not have both females and males sampled. In total, we obtained data on the plumage coloration of females and males of 876 species, representing 70% of the *ca.* 1270 species of Furnariida and Tyrannida present in the current phylogenetic hypothesis of the group (Harvey et al., 2020).

We used the R package *pavo* (Maia et al., 2013) to model plumage color in the avian tetrahedral color space (Stoddard & Prum, 2008) under the avian V system, because current evidence indicates that Tyranni passerines are not sensitive to light in the UV range (Ödeen et al., 2011). In *pavo*, we obtained the following color descriptors: relative color volume (color diversity), mean color span (color contrast), mean and maximum saturation (color vividness) (Maia et al., 2013; Stoddard & Prum, 2008). Next, we estimated plumage ornamentation for females and males using a Phylogenetic Principal Component Analysis (phyloPCA) with the phylogenetic covariance matrix and a Pagel’s lambda model using the R package *phytools* (Revell, 2009; Shultz & Burns, 2017) with scaled values of the color descriptors. We calculated the phyloPCA separately for females and males because available implementations of this method do not support phylogenetic repeated measures, i.e., a value for each sex within each species (Revell, 2009, 2012). To ease interpretation, we reversed the signs of principal component scores when necessary so that higher values of principal components corresponded to higher values of the color descriptors (i.e., higher plumage ornamentation; Table 1).

**Table 1.** Loadings of the first two principal components (PCs) of the phyloPCA of female and male plumage color. The phyloPCA was calculated separately for females and males, hence, the proportion of variance explained by each PC is also separate for each sex.

| Color Descriptors | Female PC1 | Female PC2 | Male PC1 | Male PC2 |
| --- | --- | --- | --- | --- |
| Relative Color Volume | 0.52 | -0.84 | 0.68 | -0.73 |
| Color Span | 0.81 | 0.03 | 0.86 | 0.33 |
| Mean Saturation | 0.78 | 0.44 | 0.66 | 0.38 |
| Maximum Saturation | 0.94 | 0.20 | 0.91 | 0.30 |
| Proportion of Variance | 0.58 | 0.27 | 0.61 | 0.26 |
| Cumulative Proportion | 0.58 | 0.85 | 0.61 | 0.87 |

Higher plumage color diversity, contrast and saturation are frequently used as estimates of ornamentation in comparative studies of birds (Beltrán et al., 2021; Cally et al., 2021; Cooney et al., 2019, 2022; Dale et al., 2015; Gomes et al., 2016; Mason et al., 2014; Shultz & Burns, 2017; Stoddard & Prum, 2008). However, color diversity, contrast and saturation may not reflect high ornamentation in all cases. For instance, in some species, eumelanic plumage with low color diversity, contrast and saturation likely indicates high ornamentation (e.g., the mostly black plumage of females and males of the Stub-tailed Antbird *Sipia berlepschi* and of the Amazonian Umbrellabird *Cephalopterus ornatus*). However, when comparing across larger numbers of species, even in clades in which melanin is the main component of the ornamentation (e.g., antbirds and ovenbirds; (Remsen, 2003; Zimmer & Isler, 2003), color diversity, contrast and saturation forming complex and contrasting light and dark patterns across plumage patches likely indicate high ornamentation (e.g. for species such as the Spotted Antbird *Hylophylax naevioides*, the Ocellated Antbird *Phaenostictus mcleannani*, and the Thorn-tailed Rayadito *Aphrastura spinicauda*). Therefore, to obtain a metric that is applicable in macroevolutionary scales, we assume that color diversity, contrast and saturation summarized as axes of the phyloPCA are proxies of ornamentation (Shultz & Burns, 2017).

### Sexual dichromatism

We estimated plumage sexual dichromatism by the extent of overlap (or lack thereof) between color volumes of females and males and the distance between the volumes’ centroids (G. Macedo et al., 2022). We calculated 3D volumes of females and males of each species delimited by the cartesian coordinates (x, y, z) of plumage patches in the avian tetrahedral color space using *hypervolume* (Blonder et al., 2018, 2025). Because cartesian coordinates were often highly correlated (r ≥ 0.70), which could bias volume calculation, we ran a non-phylogenetic PCA within each species and sex and used scores from the 3 resulting orthogonal principal component axes to calculate the volumes (Blonder et al., 2018). We calculated volumes with a Support Vector Machine algorithm with default values of γ = 0.5 (smoothing parameter) and ν = 0.01 (error rate parameter; Blonder et al. 2018; 2025). Briefly, *hypervolume* uses a Monte Carlo approach to randomly sample points in the parameter space.Then, an algorithm (Support Vector Machine in this case) classifies these points as being in or out of the female and male volumes, whose shapes are determined by the PCA-transformed coordinates of plumage patches. The SVM algorithm generates a tight wrap around data points, being robust against outliers and small sample sizes (Blonder et al., 2014; G. Macedo et al., 2022). Next, we calculated in *hypervolume* the Sorensen similarity index between the female and male volumes as well as the distance between their centroids. The Sorensen similarity index varies from 0 (when there is no overlap between volumes) to 1 (when there is complete overlap) and is defined as: (2 × volume of the intersection) / (female volume + male volume) (Blonder et al., 2025). We defined sexual dichromatism as: (1 – Sorensen similarity index) + (centroid distance), wherein low values correspond to species in which the sexes are more similar, and high values correspond to species in which the sexes are more different in plumage color. We added the centroid distance to obtain information on how distant (i.e., different in the color space) female and male volumes are, even when they do not overlap.

### Proxies of social competition

To develop proxies for sociosexual selection that could influence plumage coloration, we obtained data on social pair bond duration (long term, short term, or no pair bond formation, i.e., solitary species), territoriality and mating systems from (Tobias et al., 2016) and (Barber et al., 2024). We classified territoriality as being absent or present and mating systems as social monogamy or polygyny (Barber et al., 2024; Tobias et al., 2016). Although pair bond duration is correlated with mating system, both socially monogamous and polygynous Tyranni species can be solitary or have long- or short-term pair bonds; hence, pair bond duration offers additional insights into social interactions of each species (Barber et al., 2024; Tobias et al., 2016). We also obtained estimates of body size and sexual size dimorphism, which may correlate with the intensity of social competition and life history traits such as territoriality (Barber et al., 2024; Cally et al., 2021; Jones & Sheard, 2023). For instance, in songbirds, larger body sizes have been associated with higher intensities of social competition and plumage color elaboration (Dale et al., 2015). In turn, male-biased sexual size dimorphism (SSD) has been associated with male-biased sexual selection (Cally et al., 2021). Female-biased SSD can also be associated with female-biased sexual selection, although other processes such as selection for display agility in males, or fecundity selection in females, can also result in female-biased SSD (Andersson, 1994; Jones & Sheard, 2023; Pincheira-Donoso & Hunt, 2017). We obtained wing length from Avonet (Tobias et al., 2022) and used it as a proxy of body size (Dale et al., 2015; Gosler et al., 1998). Wing length is one of the best linear predictors of body size, and is widely adopted in comparative studies (Cally et al., 2021; Dale et al., 2015; Gosler et al., 1998; Odom et al., 2025; Tobias et al., 2022). We calculated SSD as the difference between log male wing length and log female wing length, with negative values corresponding to female-biased SSD (i.e., females are larger than males) and positive values corresponding to male-biased SSD (i.e., males are larger than females; (Dale et al., 2015).

Migratory behavior is associated with vulnerability to predation, with migratory species facing stronger selection for predator avoidance during migration and stopovers (Alerstam et al., 2003; Sabal et al., 2021). Migration may also be associated with stronger social competition for mates and territories, particularly in males of temperate species, because individuals must acquire territories and attract mates in short periods of time relative to non-migratory species (Badyaev & Hill, 2003; Fitzpatrick, 1997). However, studies have also suggested that year-round territoriality (i.e., in non-migratory taxa) may be associated with strong social competition in females and males (Dale et al., 2015; Odom et al., 2014). Therefore, migratory behavior (or lack thereof) and territoriality may together affect ornamentation. We used data from (Dale et al., 2015) and classified migratory behavior as present or absent.

Parental care may also influence sociosexual selection pressures, because it can bias operational sex ratios in the direction of the sex not providing parental care and increase competition for mates (Trivers, 1972; West-Eberhard, 1983; Dale et al., 2015; Drury et al., 2024). However, socially monogamous species with biparental care — which are the majority of birds (Barber et al., 2024; Dale et al., 2015) — may also face strong competition for mates and other resources (Jones & Montgomerie, 1991; Kokko & Johnstone, 2002; Kraaijeveld, 2003; Kraaijeveld et al., 2007; Odom et al., 2014). We obtained data on parental care from the primary literature and species accounts of the Birds of the World (BOW; Billerman et al., 2025). We classified parental care as biparental or female-only, because in Tyranni there is currently no evidence of species with exclusive or frequent male-only parental care (Billerman et al., 2025). As another proxy of parental investment, we obtained clutch sizes (Jetz et al., 2008; Ricklefs, 2010) from BOW (Billerman et al., 2025), considering mean values reported for each species. We acknowledge that clutch size by itself is a limited proxy of parental investment as egg volume interacts with clutch size in different reproductive strategies (Fargevieille et al., 2023); however, information on egg volume is not currently available for most Tyranni species.

### Environmental factors and proxies of natural selection

To develop proxies for natural selection pressures that could shape plumage coloration, we considered habitat type, locomotion style and nest type, which are associated with camouflage and predator avoidance (Davidson et al., 2017; Marcondes & Brumfield, 2019; Tobias & Pigot, 2019). In birds, species living in darker habitats tend to show darker plumage, and species living in lighter habitats tend to show lighter plumage, which is thought to favor camouflage against predators (Beco et al., 2021; Endler, 1992; Marcondes & Brumfield, 2019). Furthermore, species with terrestrial lifestyles and/or with open nest types (e.g. open cups) are thought to be more exposed to terrestrial and aerial predators than species with insessorial (perching) habits and/or closed nest types (e.g., domes and cavities; (Campos et al., 2009; Davidson et al., 2017; Lima, 2009; Tobias & Pigot, 2019). We obtained data on habitat type and locomotion from Avonet (Tobias et al., 2022). We classified habitat type as closed (forests and woodlands) and open habitats (e.g., savannas, shrublands, grasslands, coastal and aquatic vegetation), and locomotion styles as terrestrial (generalist and cursorial species moving mostly on the ground) and insessorial. Based on primary literature and species accounts of BOW (Billerman et al., 2025), we classified nest types as open cups, domes (closed nest structures such as oven-like nests of ovenbirds), primary cavities (i.e., excavated by individuals of the focal species themselves) and secondary cavities (i.e., naturally occurring cavities such as tree hollows or cavities built by other species such as mammals or woodpeckers).

Climate and climatic seasonality affect habitat structure, resource availability and social competition (G. Macedo et al., 2022; R. H. Macedo & Machado, 2013). For instance, more seasonal environments have been associated with stronger competition for territories and mates, which may result in higher ornamentation (Badyaev & Hill, 2003; R. H. Macedo & Machado, 2013). Additionally, temperature and precipitation may also directly affect plumage coloration, e.g., via thermal melanism and the Gloger’s rule (G. Macedo et al., 2024; Marcondes et al., 2021). Therefore, we obtained annual mean temperature (°C), annual mean precipitation (mm) and temperature seasonality (coefficient of variation; %) from (Harvey et al., 2020) (2020), who used species distribution maps and data from WorldClim (Hijmans et al., 2005). As a proxy for energy and resource availability in food webs, which also influences social competition, we used estimates of net primary productivity from Cally et al., (2021), which is based on remote sensing data of photosynthetic activity by (Zhao et al., 2005).

### Statistical Analyses

We conducted all statistical analyses in R version 4.5.0. To account for the phylogenetic non-independence of species, we used the phylogeny of Tyranni passerines proposed by (Harvey et al., 2020) and specified the inverse phylogenetic covariance matrix as a group-level effect in *brms* (Bürkner, 2017; Bürkner et al., 2024). We tested sexual dichromatism, female and male PC1 and PC2 as response variables. In the models with PCs, we specified female and male PC1, and, in a separate model, female and male PC2 as response variables in phylogenetic multi-response models (Halliwell et al., 2025) to investigate the evolutionary drivers of female and male plumage color elaboration while accounting for intercorrelations that may arise due to shared genetic/developmental backgrounds between sexes (Lande, 1980). We applied the Yeo-Johsnson transformation to PC1 and PC2 of females and males because these variables have both positive and negative values and had high skewness (Yeo & Johnson, 2000). As predictors, we specified social pair bond duration, mating system, sexual size dimorphism, wing length, territoriality, parental care, clutch size, locomotion style, habitat type, nest type, net primary productivity, and temperature, precipitation, and temperature seasonality. We log-transformed positive continuous variables (clutch size, wing length, net primary productivity, and temperature seasonality) and scaled sexual size dimorphism (which is already a difference of logs) to a mean of 0 and standard deviation of 1.

We also specified two-way interactions between predictors that we expect to simultaneously affect female and male plumage elaboration. We tested the interaction between territoriality and migration, because migrating territorial species are predicted to face strong social competition for mates and territories in breeding grounds, particularly in males (Badyaev & Hill, 2003). On the other hand, species that defend territories year-round (i.e., are non-migratory) may also face strong social competition over territories (Tobias et al., 2016). We also tested the interaction between habitat type and migration, because migratory species inhabiting open habitats may be more prone to predation, which may favor cryptic plumage coloration (Alerstam et al., 2003; Simpson et al., 2015). Additionally, we tested the interaction between nest type and parental care, because we expect that males are affected by nest type only when they provide parental care. Cavity nests also provide protection against visual predators during incubation, and secondary cavity nests have been associated with greater territorial aggression, which may be associated with higher plumage elaboration (Lipshutz et al., 2025; Lipshutz & Rosvall, 2021). As mating systems are often accompanied by other traits that may affect female and male ornamentation (Barber et al., 2024; Dale et al., 2015), we tested two-way interactions between mating system and territoriality, mating system and wing length (body size), and mating system and sexual size dimorphism. Lastly, as seasonality can strengthen social competition for resources such as territories even in non-migratory species (Badyaev & Hill, 2003; R. H. Macedo & Machado, 2013), we tested a two-way interaction between territoriality and temperature seasonality. Due to the large number of predictors and interactions, we ran models with a single or two predictors (when testing a two-way interaction) as well as a full model with all predictors and interactions to assess if conclusions would differ depending on the variables considered (Dale et al., 2015). Results of the full model were qualitatively similar to uni- and bivariate models; thus, we only report the results of the full model. We ran Bayesian models in 4 Markov chains for 10,000 iterations (50% as warm-up) and specified a thinning rate of 1 (Link & Eaton, 2012). We adopted regularizing priors (Lemoine, 2019) and considered that an effect was likely to be present when 95% credible intervals did not include zero.

To obtain post-hoc contrasts between females and males of the PCA axes of color variation, we compared PC1 and PC2 scores between females and males with phylogenetic repeated measures models with sex added to the predictors, and species names as a group-level effect in addition to the inverse phylogenetic matrix (Paltrinieri et al., 2025). We calculated post-hoc contrasts across the sexes and categorical predictors with the R package *emmeans* (Lenth et al., 2025). Because species of Tyranni nesting in primary cavities have biparental care but lack female-only parental care, there are cases of zero-valued observations when testing the interaction between nest type and parental care, which could affect the models’ estimates. Therefore, to account for unbalanced sample sizes across categorical predictors as well as ignore zero-valued observations, we used “cells*”* weights, which assign weights to marginal means according to the frequencies of observations while ignoring zero-valued observations (Lenth et al., 2025).

We imputed missing values in predictors using the R package *Trait Data Imputation with Phylogeny TDIP* (Gendre et al., 2024), as 67 species had missing values of female or male wing length, 140 species had missing values of nest type, 272 species has missing values of clutch size, 117 had missing values of parental care, and 88 species had missing values of net primary productivity, temperature, precipitation and temperature seasonality. We employed phylogenetic missForest, multinomial logistic regression, and k-nearest neighbor algorithms, and used hard voting to obtain a majority prediction across algorithms (Gendre et al., 2024). To assess the sensitivity of our analyses to imputation, we also ran models considering only the species that had complete data for all predictors (467 species; 53% of the species in this study).

## Results

### Plumage ornamentation and sexual dichromatism

We observed the lowest values of plumage sexual dichromatism in the ovenbirds (Furnariidae), which is a clade mostly comprised of sexually monochromatic species, such as the Azara’s Spinetail (*Synallaxis azarae*). Nonetheless, there is large variation in sexual dichromatism across Tyranni, and other clades also have species with low sexual dichromatism such as the Russet Antshrike (*Thamnistes anabatinus*). Along the continuum, we observed intermediate to high values of sexual dichromatism in species of antbirds (Thamnophiliade), manakins (Pipridae) and cotingas (Cotingidae), such as the Common Scale-backed Antbird (*Willisornis poecilinotus*), Black-necked Red-Cotinga (*Phoenicircus nigricollis*), and the Swallow-tailed Manakin (*Chiroxiphia caudata*) (Figure 1).

**Figure 1.**
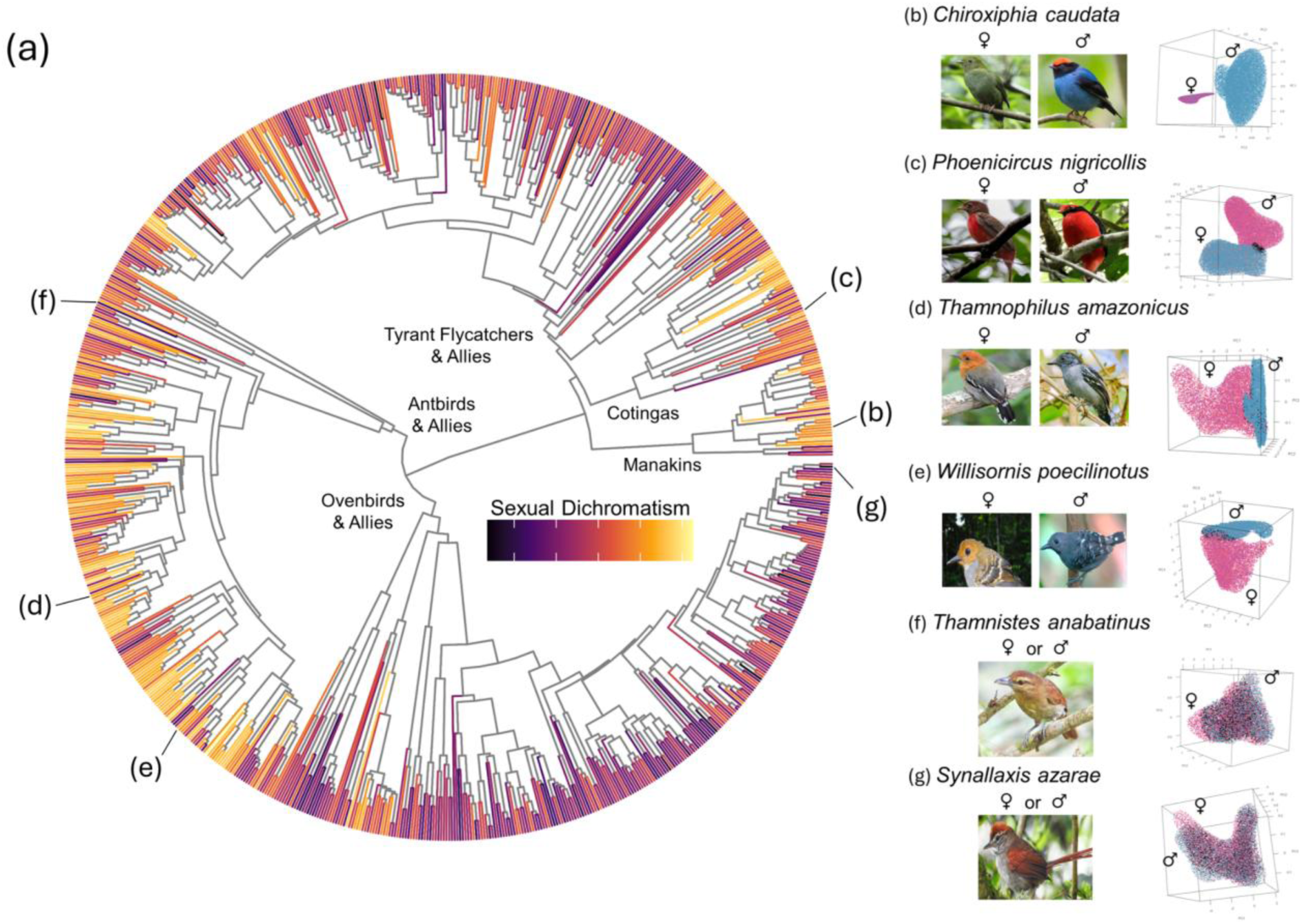
Phylogeny of Tyranni passerines showing the 876 species included in the analysis (a). Terminal branches of the tree are colored according to sexual dichromatism obtained with 3D volumes. Photographs show examples of Tyranni species, their position in the phylogeny, and their respective female (in pink) and male (in blue) color volumes (when present, overlap between volumes is represented in black; b-g). b and c are examples of species with male-biased sexual dichromatism. d and e are examples of species with female-biased sexual dichromatism. f and g are examples of species with inconspicuous and elaborate monomorphisms, respectively. Credits of photographs and Creative Commons licenses: b, female: Jean-Paul Boerekamps (CC0); b, male: Carlos Moura (CC-BY-NC); c, female: Hector Bottai (CC-BY-SA); c, male: Nick Athanas (CC-BY-SA); d, female: Edwin Múnera Chavarría (CC-BY-NC); d, male: Anderson Sandro (CC-BY-NC); e, female: Rafael Homobono Naiff (CC-BY-NC); e, male: Mike Melton (CC-BY-NC); f: Nelson Apolo (CC-BY-NC); g: Saul Arias (CC-BY-NC).

Phylogenetic principal components had overall similar loadings in females and males. In both sexes, PC1 corresponds to an axis of plumage color complexity, with increasing values of color diversity (relative color volume), contrast (span) and vividness (saturation), whereas PC2 mostly corresponds to an axis of increasing vividness but decreasing color diversity (Table 1).

### Proxies of social competition are associated with higher female and male ornamentation and lower sexual dichromatism

We found that social pair bond duration predicts female and male ornamentation as well as sexual dichromatism (Table S1-S3). In species with long-term social pair bonds, females are slightly more ornamented than males (PC1: β = 0.16 [0.07 : 0.24]; henceforth, numbers in square brackets are 95% CI; Figure 2). In contrast, males are more ornamented than females in species with short-term pair bonds (PC1: β = −0.43 [−0.62 : −0.24]) and in solitary species (PC1: β = −0.56 [−0.85 : - 0.27]; Table S4). Sexual dichromatism is highest in solitary species (Figure 2).

**Figure 2.**
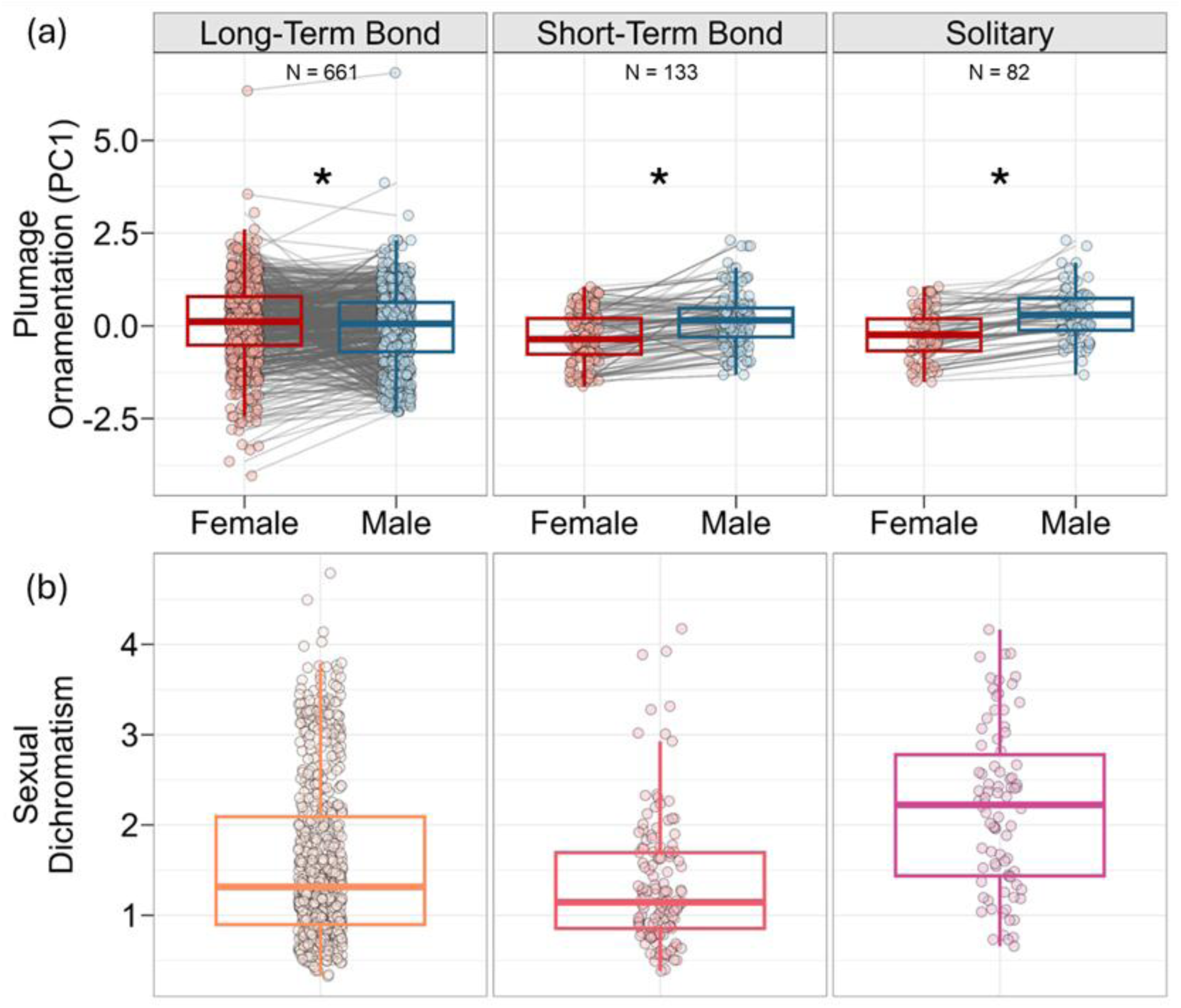
Species with long-term social pair bonds show higher female than male ornamentation, whereas those with short-term pair bonds and solitary species show higher male than female ornamentation. Boxplots are in the style of Tukey and show (a) PC1 of females and males and (b) sexual dichromatism in each social pair bond category. PC1 was obtained with a phylogenetic PCA and represents an axis of increasing plumage color diversity, contrast and saturation in females and males. Sexual dichromatism was calculated with 3D volumes and represents how different females and males are in the avian tetracolor space. Asterisks indicate comparisons that are credibly different from zero. Lines connect points referring to female and male plumage ornamentation of the same species. Numbers in the upper plots are sample sizes in each category.

We tested the interaction between territoriality and social mating system because in socially monogamous Tyranni species, both sexes often defend territories, which may favor elaborate monomorphism. As expected, we found that the interaction between territoriality and social mating system predicts female and male plumage ornamentation (Tables S1 and S2). Females and males of territorial and socially monogamous species are similarly ornamented (PC1: β = 0.06 [−0.02 : 0.14]), and, correspondingly, show low sexual dichromatism (Figure 2). In contrast, in non-territorial and socially monogamous species, females are more ornamented than males (PC1: β = 0.58 [0.04 : 1.11]). In species with socially polygynous mating systems, females and males are similarly ornamented in territorial species (PC1: β = −0.34 [−0.74 : 0.07]), but females are less ornamented than males in non-territorial species (PC1: β = −0.58 [−0.86 : −0.30]). Correspondingly, non-territorial polygynous species show high levels of sexual dichromatism (Figure 3).

**Figure 3.**
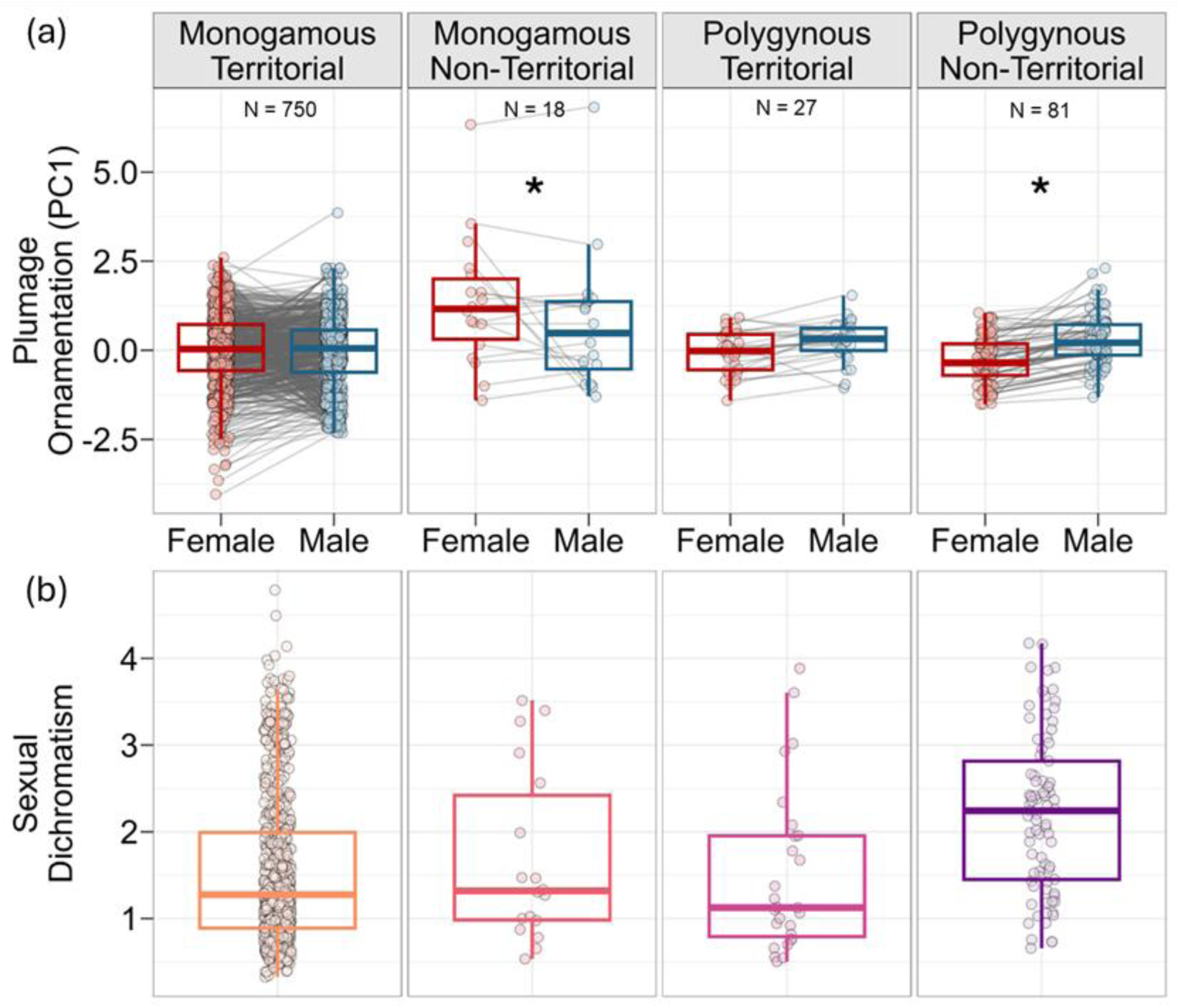
Territoriality and social monogamy are associated with more similar ornamentation between the sexes and lower sexual dichromatism. Boxplots are in the style of Tukey and show (a) PC1 of females and males and (b) sexual dichromatism of the interaction between social mating systems and territoriality. PC1 was obtained with a phylogenetic PCA and represents an axis of increasing plumage color diversity, contrast and saturation in females and males. Sexual dichromatism was calculated with 3D volumes and represents how different females and males are in the avian tetracolor space. Asterisks indicate comparisons that are credibly different from zero. Lines connect points referring to female and male plumage ornamentation of the same species. Numbers in the upper plots are sample sizes in each category.

The interaction between territoriality and migratory behavior, which may affect sociosexual selection pressures (e.g., competition for mates and territories on breeding grounds) predicted female ornamentation, male ornamentation, and sexual dichromatism (Tables S1-S3). Females are more ornamented than males in territorial and non-migratory species (PC1: β = 0.13 [0.04 : 0.20]), whereas in non-territorial and migratory species females and males show similar ornamentation (PC1: β = −0.14 [−0.57 : 0.89]), although sample sizes of non-territorial and migratory species are small (N = 8; Figure 4). Conversely, males show greater ornamentation than females and sexual dichromatism is higher in migratory and territorial (PC1: - 0.49 [−0.73 : −0.26]), as well as in non-migratory and non-territorial species (PC1: - 0.41 [−0.70 : −0.13]; Figure 4; Table S4).

**Figure 4.**
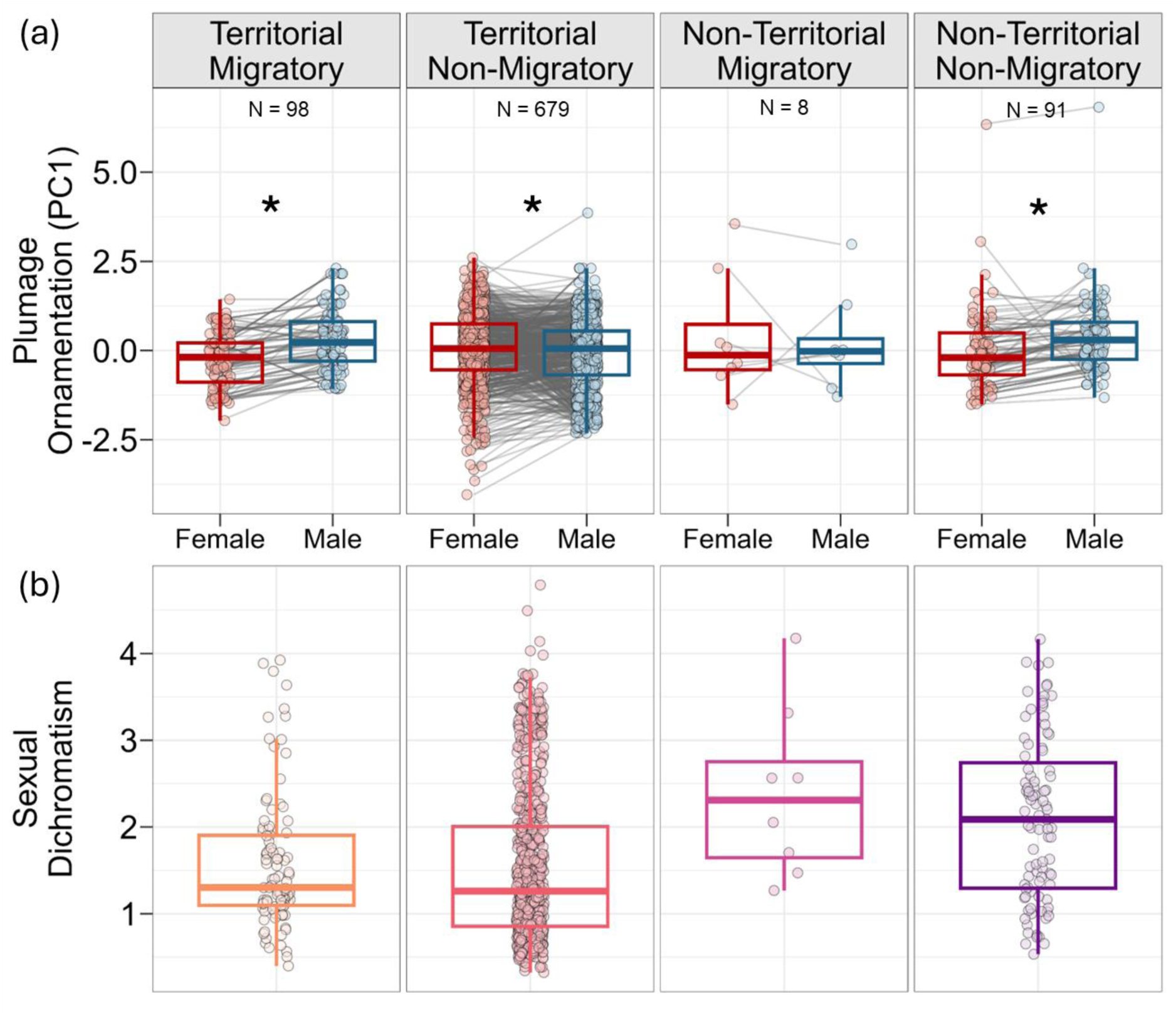
Territoriality and migratory behavior, as well as lack of territoriality and migratory behavior, are associated with greater ornamentation in males than in females, whereas territoriality and lack of migratory behavior are associated with greater ornamentation in females than in males. Boxplots are in the style of Tukey and show (a) PC1 of females and males and (b) sexual dichromatism of the interaction between territoriality and migratory behavior. PC1 was obtained with a phylogenetic PCA and represents an axis of increasing plumage color diversity, contrast and saturation in females and males. Sexual dichromatism was calculated with 3D volumes and represents how different females and males are in the avian tetracolor space. Asterisks indicate comparisons that are credibly different from zero. Lines connect points referring to female and male plumage ornamentation of the same species. Numbers in the upper plots are sample sizes in each category.

Because climatic seasonality may constrain available time and resources to defend territories, we tested the interaction between territoriality and temperature seasonality. Supporting this prediction, we found that temperature seasonality interacts with territoriality to explain male ornamentation (PC1: β = 0.23 [0.04 : 0.42]); however, in females, this effect was not credibly different from zero (PC1: β = 0.13 [−0.05 : 0.31]). In territorial species, higher temperature seasonality is associated with higher male ornamentation, whereas in non-territorial species ornamentation is independent of temperature seasonality (Figure S2; Table S2). Net primary productivity, which is also a proxy of environmental energy and resource availability, showed non-credible negative effects on female ornamentation (PC2: β = −0.14 [−0.30 : 0.01]) and non-credible positive effects on male ornamentation (PC1: β = 0.11 [−0.01 : 0.22]).

### Lower vulnerability to predation and biparental care are associated with higher female and male ornamentation and lower sexual dichromatism

We tested an interaction between parental care and nest type because we expected that males would only be exposed to predators during nesting when they provide parental care. We found that parental care and nest type predict female and male ornamentation. In species that nest in secondary cavities, males are more ornamented than females in species with biparental care (PC1: β = −0.24 [−0.41 : - 0.07]), but females and males show similar ornamentation in species with female-only parental care (PC1: β = −0.13 [−0.75 : 0.48]; however, the sample size in this category is small; N= 12; Figure 5). Likewise, females and males are similarly ornamented in species that nest in domes with biparental care (PC1: β = −0.10 [−0.25 : 0.06]), and species that nest in primary cavities (PC1: β = −0.24 [−0.54 : 0.04]; in Tyranni, these species exclusively show biparental care). In contrast, in species that nest in domes and show female-only parental care, males are more ornamented than females (PC1: β = −0.61 [−1.17 : −0.06]). Species that nest in secondary and primary cavities showed the lowest levels of sexual dichromatism (Figure 5b). In contrast, males are more ornamented than females and sexual dichromatism is higher in species with female-only parental care that nest in open cups (PC1: β = - 0.59 [−0.87 : −0.32]; Figure 4). Unexpectedly, we found that females show greater ornamentation than males in species with biparental care that nest in open cups (PC1: β = 0.36 [0.25 : 0.47]; Figure 5; Table S4).

**Figure 5.**
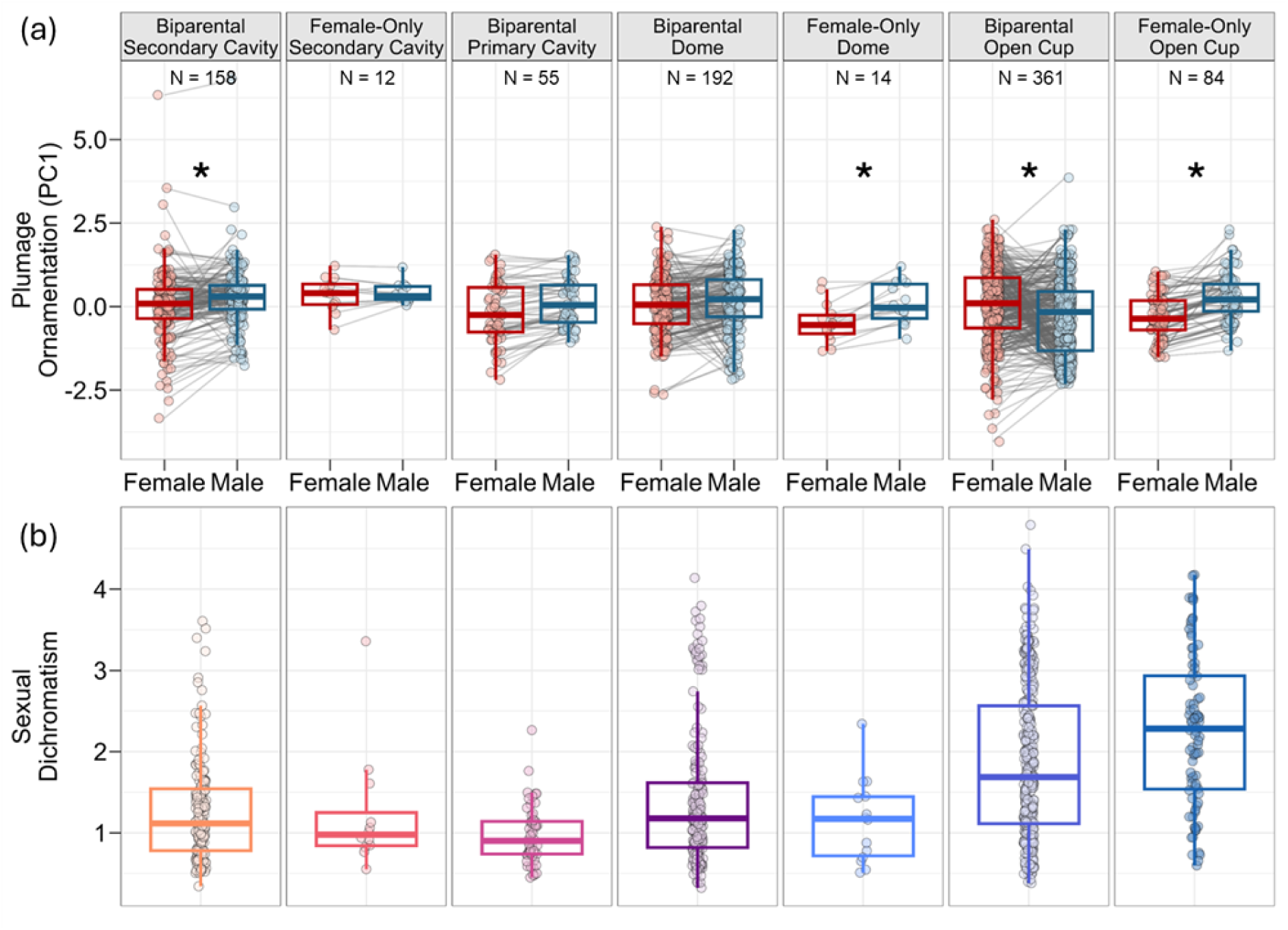
Biparental care and closed nest types (secondary/natural and primary/excavated cavities and domes) are associated with more similar ornamentation between the sexes and lower sexual dichromatism, whereas female-only parental care in open cup nests are associated with higher male ornamentation and higher sexual dichromatism. In contrast, biparental care and open cup nests are associated with higher female ornamentation. Boxplots are in the style of Tukey and show (a) PC1 of females and males and (b) sexual dichromatism of the interaction between parental care and nest type. PC1 was obtained with a phylogenetic PCA and represents an axis of increasing plumage color diversity, contrast and saturation in females and males. Sexual dichromatism was calculated with 3D volumes and represents how different females and males are in the avian tetracolor space. Asterisks indicate comparisons that are credibly different from zero. Lines connect points referring to female and male plumage ornamentation of the same species. Numbers in the lower right corners of plots are sample sizes in each category.

Migratory species living in open habitats are likely more vulnerable to predators than species living in closed habitats and non-migratory species. Therefore, we tested the interaction between habitat type and migration. We found that, for species living in closed habitats, females and males show similar ornamentation in migratory species (PC1: β = −0.26 [−0.71 : 0.19]), and females are more ornamented than males in non-migratory species (PC1: β = 0.17 [0.07 : 0.27]). Conversely, in open habitats, females show lower ornamentation than males in migratory species (PC1: β = −0.50 [−0.74 : −0.24]) and non-migratory species (PC1: β = −0.20 [−0.36 : −0.05]; Figure S1). Migratory species showed the highest levels of sexual dichromatism in relation to non-migratory species (Figure S1). In relation to locomotion styles, we found that females and males show similar ornamentation in terrestrial species (PC1: β = 0 [−0.08 : 0.09]), and insessorial species (PC1: β = −0.01 [−0.18 : 0.16]; Figure S3; Table S4).

Contrasts with PC2 values of females and males were similar to contrasts with PC1 (Table S5). Furthermore, the models without imputation of missing data in predictors showed generally similar results to the models with imputation. However, instances in which females showed greater ornamentation than males were no longer credibly different from zero. That is, when removing imputed missing data, females and males tended to show similar ornamentation in species with long term social pair bonds, in non-migratory territorial species, non-migratory species living in closed habitats, and non-territorial monogamous species (Supplementary Material *Comparisons between the models with and without imputation*; Tables S6-S10).

### Female-biased parental investment and male-biased sexual size dimorphism (SSD) are associated with lower female and higher male ornamentation

Biparental or female-only parental care may interact with clutch size to determine parental investment, affecting sociosexual selection pressures. Supporting this expectation, we found that the interaction between clutch size and parental care predicts female and male ornamentation. Males show higher ornamentation than females in species with female-only parental care as clutch size increases (PC1: β = −0.53 [−0.78 : −0.29]; Figure S4), but, in species with biparental care, clutch size does not credibly affect female and male ornamentation (PC1: β = 0.08 [0 : 0.15]; Figure S4).

Lastly, we tested the interaction between social mating systems and SSD because male-biased SSD is often associated with polygynous species. Consistently with this expectation, we found that higher male ornamentation is associated with male-biased SSD , but only in species with polygynous social mating systems (PC2: β = 0.15 [0.03 : 0.27]). In socially monogamous species, SSD has no effect on male ornamentation (Figure 6). Female ornamentation was not credibly associated with SSD (PC2: β = 0.11 [−0.02 : 0.24]; Table S3).

**Figure 6.**
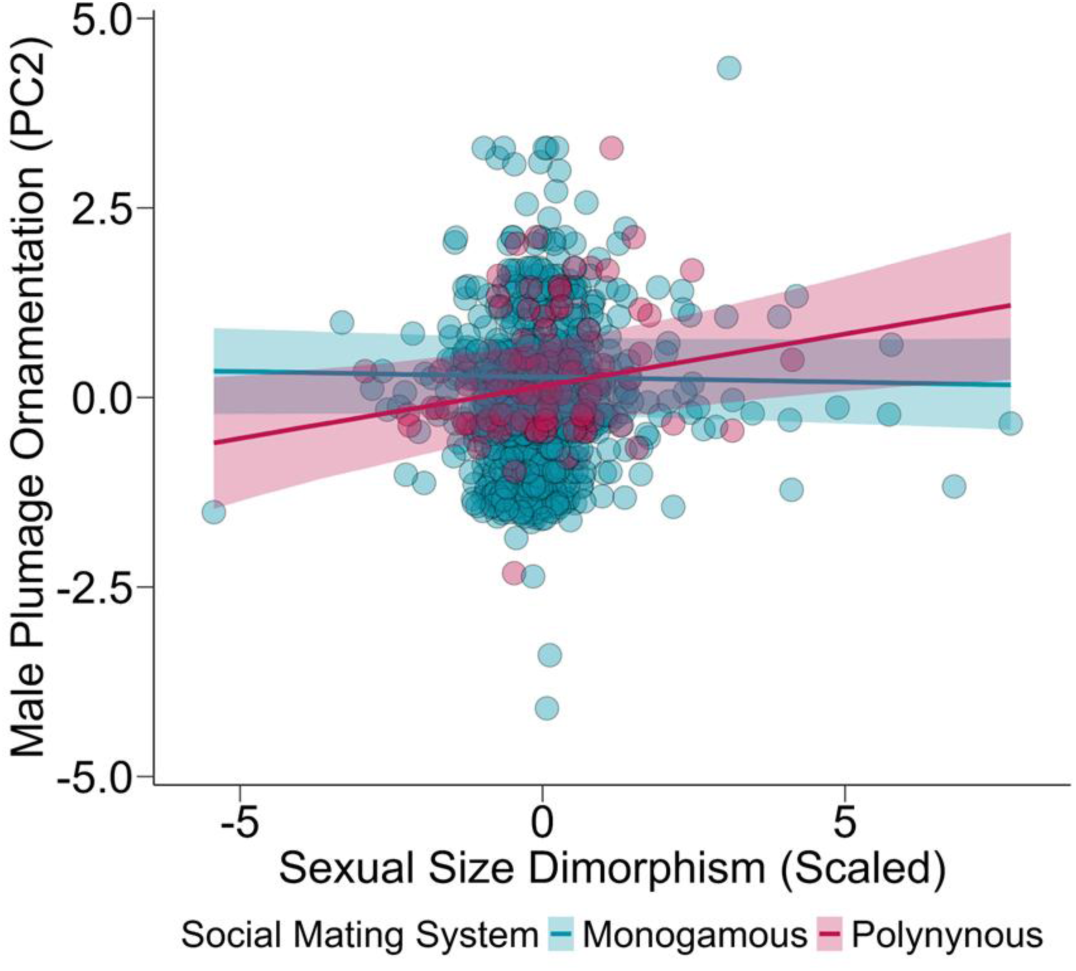
Male-biased sexual size dimorphism (positive values in the x axis) is associated with higher male ornamentation in socially polygynous species, but not in socially monogamous species. PC1 was obtained with a phylogenetic PCA and represents an axis of increasing plumage color diversity, contrast and saturation in females and males. Each point represents a species, which are colored according to territoriality and social mating system categories. Shaded areas are 95% CI.

## Discussion

Using plumage ornamentation of Tyranni as a model system for the evolution of social signaling traits and statistically controlling for correlations between females and males, we showed that higher ornamentation in both females and males is associated with proxies of sociosexual selection pressures. Notably, similarly high ornamentation and the corresponding lower sexual dichromatism (i.e., elaborate monomorphism) are more prevalent than male-biased sexual dichromatism in Tyranni. Elaborate monomorphism was associated with similar sociosexual selection pressures between the sexes, such as in species with socially monogamous mating systems, biparental care, and territoriality. In contrast, we found that lower ornamentation in females relative to males is associated with higher female parental investment (female-only parental care and larger clutch sizes), higher vulnerability to predation such as in migratory species living in open habitats, as well as higher climatic seasonality. We also observed instances in which females are more ornamented than males, such as in species with long-term pair bonds, and in territorial and non-migratory species living in closed habitats. Other factors associated with social competition and vulnerability to predation such as nest type and parental care also affected female and male ornamentation. Primary/excavated cavities and domes in species with biparental care, and secondary cavities in species with female-only parental care were associated with similarly high female and male ornamentation. In contrast, open-cup nests in species with female-only parental care, and secondary/natural cavities in species with biparental care were associated with higher ornamentation in males than in females, whereas open-cup nests were associated with higher ornamentation in females than in males in species with biparental care. Locomotion styles and net primary productivity did not consistently affect female and male ornamentation. Lastly, we found that male-biased sexual size dimorphism (SSD) is associated with higher male ornamentation in polygynous species, but female ornamentation was not associated with SSD. Below, we discuss how natural and sociosexual selection pressures shape ornamentation in females and males.

### Sociosexual selection shapes female and male plumage ornamentation

Life history traits associated with sociosexual selection revealed key circumstances in which females and males show similar ornamentation in Tyranni. These similarities ranged from similarly low (inconspicuous monomorphism) to similarly high ornamentation (elaborate monomorphism) in females and males. In socially monogamous and territorial species, both females and males show similar ornamentation, with many cases corresponding to elaborate monomorphism. Studies investigating social competition have shown that stronger territorial responses are associated with higher elaboration of social traits in both sexes. For instance, playback experiments comparing species of antbirds (G. Macedo et al., 2021), Neotropical wrens (Mennill, 2006), and shorebirds (Lipshutz, 2017) have shown that higher plumage ornamentation, song duet displays, body and weapon sizes, as well as dry seasons in relation to wet seasons (Fedy & Stutchbury, 2005) are associated with stronger aggressive displays in both sexes. Likewise, species that engage in complex social behaviors such as mixed species flocks (Beco et al., 2021) and cooperative breeding (Rubenstein & Lovette, 2009), in which females and males show similar “sex roles” with more similar parental investment (Fargevieille et al., 2023; Odom et al., 2021, 2025) tend to show higher female and male ornamentation and lower sexual dimorphism. Furthermore, within and across species of birds, both female and male ornamentation are positively associated with proxies of body condition (e.g., body size and immunity) and fitness (e.g., number and survival of offspring) (Nolazco et al., 2022). The present study adds to this body of evidence showing that, just as male social traits, female social traits may also evolve due to direct sociosexual selection pressures. Importantly, our results provide support for predictions of (West-Eberhard, 1979, 1983, 2014), indicating that when the sexes engage in social competition with similar intensities, social traits show sexually monomorphic elaboration.

Interestingly, we observed species with female-biased sexual dichromatism. For instance, we found that females show higher ornamentation than males in species with long-term pair bonds, with biparental care that nest in open cups, or that are non-migratory and territorial and live in closed habitats. However, these cases no longer showed credible effects in the models removing species with missing data in predictions, and, thus, should be interpreted with care. Examples of species with female-biased sexual dichromatism are most notable in antbirds, such as the Large-billed Antwren (*Herpsilochmus longirostris*), the Xingu Scale-backed Antbird (*Willisornis vidua*), and the Pacific Antwren (*Myrmotherula pacifica*) (Zimmer & Isler, 2003). High female ornamentation in species nesting in open cups seems unexpected due to predation risks. Nonetheless, available evidence indicates that some female antbirds incubate eggs mostly at night (Zimmer & Isler, 2003), which could protect them from visual diurnal predators and possibly explain their higher ornamentation. However, very few Neotropical species such as Tyranni passerines have the natural history and behavioral ecology data necessary to investigate the drivers of higher ornamentation in females in relation to males (Winkler et al., 2020; Zimmer & Isler, 2003). Field-based research, especially in the tropics, is indispensable to formulate and test hypotheses about the ecological and evolutionary processes shaping social competition and ornamentation in females and males (Soga & Gaston, 2025; Tobias et al., 2012).

We also observed species in which females are less ornamented than males. Remarkably, these instances of male-biased sexual dichromatism, which have been emphasized in studies investigating sociosexual traits (Ah-King, 2022), are less common than elaborate monomorphism in Tyranni. The historical emphasis on male-biased trait elaboration is perhaps related to the bias in studying birds from temperate zones (Ah-King, 2022; Riebel et al., 2019). This possibility is supported by other studies that found that male-biased sexual dimorphism in plumage and songs is more common in temperate songbirds, whereas females and males of tropical species tend to be more similarly ornamented (Dale et al., 2015; Odom et al., 2025). In Tyranni, lower female ornamentation in relation to males is associated with polygynous species that lack territoriality, such as the Pin-tailed Manakin (*Ilicura militaris*), the Swallow-tailed Manakin (*Chiroxiphia caudata*) and the Andean Cock-of-the-rock (*Rupicola peruvianus*), or that are both migratory and territorial, such as the Vermilion Flycatcher (*Pyrocephalus rubinus*). Likewise, we observed lower female ornamentation in species with higher female investment in parental care, such as in species with female-only parental care and with larger clutch sizes. Moreover, higher climatic seasonality (a proxy for resource variability) in territorial species, male-biased sexual size dimorphism in polygynous species were associated with higher male ornamentation, but female ornamentation was not consistently associated with climatic seasonality nor female-biased sexual size dimorphism. Net primary productivity, which is another factor associated with resource availability, did not show consistent effects on the ornamentation of either sex, with only a weak trend of increasing male ornamentation and decreasing female ornamentation with increasing net primary productivity. Overall, these results also corroborate predictions of sociosexual selection, whereby stronger social competition acting on a given sex (in this case, male-male competition and female mate choice acting on male plumage), leads to sex-biased elaboration of social traits (Darwin, 1871; West-Eberhard, 1983).

Previous studies conducting taxonomic family-level analyses found that intermediate levels of climatic seasonality are associated with higher ornamentation in females and males of birds (López-Idiáquez et al., 2025; G. Macedo et al., 2022), whereas the present study and others with broader taxonomic scope find that males are more affected by climatic seasonality than females (Barber et al., 2024; Cooney et al., 2022), suggesting that the positive effect of seasonality on ornamentation is generally more common in males than in females. Additionally, because female ornamentation was not associated with sexual size dimorphism, body size may be more relevant to mediate social competition in males than in females. In a study investigating all passerines, (Cally et al., 2021) found that male-biased sexual size dimorphism, but not sexual dichromatism, is associated with higher speciation rates in songbirds, another key prediction of sociosexual selection theory (West-Eberhard, 1983). Other studies across diverse taxa have also reported lack of associations between male-biased sexual dichromatism and proxies of sociosexual selection (Huang & Rabosky, 2014; Kraaijeveld et al., 2011). Our study helps to explain these findings by showing that lower sexual dichromatism can also evolve due to similarly high social-sexual selection pressures in females and males, and not solely from low levels of sociosexual selection (Kraaijeveld et al., 2007). Specifically, our study shows that not all monomorphisms evolve equally, and that distinguishing between elaborate and inconspicuous monomorphism, in addition to sexual dimorphism, is necessary to understanding the effects of sociosexual selection on trait evolution.

### Trade-offs between natural and sociosexual selection in females and males

Previous studies have shown that higher ornamentation increases predation risk (Godin & McDonough, 2003; Stuart-Fox et al., 2003). Migration is also associated with increased predation risk as well as strengthened competition for mates and ecological resources such as food and territories (Alerstam et al., 2003; Tobias & Pigot, 2019). Furthermore, some evidence indicates that migrating females may be more profitable prey items due to their higher investment in energetic/fat reserves during egg production, lower agility before egg laying, and greater exposure in the nest in the breeding grounds of species with female-only parental care (Lima, 2009; Magnhagen, 1991). We found that females of migratory species living in open habitats show lower ornamentation than males. Our results corroborate other studies investigating female and male ornamentation in relation to migration or habitat type (Medina et al., 2017; Simpson et al., 2015). For instance, in wood warblers (Parulidae), female, but not male ornamentation decreases with increasing migration distances and breeding latitudes (Simpson et al., 2015). Likewise, in fairy-wrens (Maluridae), which are not migratory, open habitats are associated with lower female, but not male ornamentation (Medina et al., 2017). The present study shows that, in Tyranni, the negative effect of migration on female ornamentation is only apparent in species living in open habitats. This pattern suggests that, in the trade-off between natural and sociosexual selection generated by migration in open habitats, females invest more in crypsis and survival, possibly because they are at greater risk of predation. In contrast, males of migratory species living in open habitats invest more in sociosexual signaling. Our findings suggest that closed habitats may provide some protection against visual predators for females of migratory species.

The trade-off between natural and sociosexual selection in female and male ornamentation was also noticeable when considering nest types and parental care. We found that females and males show similarly high ornamentation in species with biparental care nesting in primary cavities and domes. Conversely, males are more ornamented than females in species with female-only parental care that nest in domes and open cups. In secondary cavities, we observed that males are more ornamented than females in species with biparental care, but females and males show similarly high ornamentation in species with female-only parental care. Secondary cavities are a limiting resource that has been associated with stronger social competition and territorial aggression in females and males (Lipshutz et al., 2025; Lipshutz & Rosvall, 2021). Our findings support this pattern suggesting that, in Tyranni, plumage ornamentation may mediate social competition for natural cavities in females and males, but in females of Tyranni this effect is more pronounced when they are the sole providers of parental care. Together, our findings suggest that the costs associated with visual predators affect female ornamentation more strongly than male ornamentation during migration and in open habitats, and when females are the sole providers of parental care in more exposed nest types (Medina et al., 2017; Simpson et al., 2015). In contrast, in species that are not migratory, live in closed habitats, show biparental care and/or nest in more protected (closed) nests, females and males show similar levels of ornamentation.

### Female-biased trait elaboration: going beyond the sex binary and other paths of future research

Sexual variation in coloration has been widely tested, especially in birds (Badyaev & Hill, 2003; Bailey, 1978; Barber et al., 2024; Cooney et al., 2022; Hamilton, 1961; Shultz & Burns, 2017). However, there is more work to be done. For instance, better knowledge of social trait variation is needed across a greater diversity of modalities, such as acoustic and chemical signals, as well as weapons (e.g, horns, antlers, spurs), and behaviors (Ah-King, 2022; Odom et al., 2025; Riebel et al., 2019). Moreover, species with female-biased trait elaboration provide an excellent opportunity to investigate the molecular basis of female ornamentation, which is poorly understood in relation to male ornamentation. For instance, studies investigating genetic and hormonal variation are relevant to elucidate whether the expression of female and male ornaments rely on similar or different molecular mechanisms (Enbody et al., 2022; Merondun et al., 2024).

Our macroevolutionary study provides a broad framework for how increased sociosexual selection shapes higher female ornamentation, but more experimental and natural history studies are necessary to better understand these socioecological factors. For example, differences in territoriality and parental care investment across sexes are poorly known for most species. Despite shared territorial defense and biparental care in many species of Tyranni, the degree to which the sexes engage in these behaviors may vary. In some species, the sexes defend territories and invest in parental care with similar intensities (Greenberg & Gradwohl, 1983; Perrella et al., 2016; Sánchez-Martínez & Londoño, 2016; Skutch, 1996), whereas in others, males show stronger territorial responses than females, and females invest more in parental care than males (Caicedo & Londoño, 2017; Fedy & Stutchbury, 2005; G. Macedo et al., 2021; Reinert, 2009; Rompré & Robinson, 2008; Skutch, 1996; Tobias et al., 2011). Additional studies connecting variation in territorial and parental investment with the degree of ornamentation will provide finer grained data to unveil sociosexual selection pressures shaping female and male traits.

Understanding variation across traits is also relevant for developing approaches to study the continuum of sex differences and similarities, beyond *a priori* binary categorization of sex (Massa et al., 2023; McLaughlin et al., 2023; Sharpe et al., 2023). To date, the most prevalent way in which traits are measured, including plumage color as in the datasets used in the present study, is by assigning sex to a specimen first and then testing whether there are statistical differences between the female and male categories. However, this approach may artificially binarize the continuous variation across traits of animals. Multivariate approaches may offer an alternative to this *a priori* binarization (McLaughlin et al., 2023). In Bayesian approaches, models that use distributional mixtures offer a promising method to model uni-, bi- and multimodal distributions of trait variation, which can then be used to infer ecological and evolutionary processes (Beraha et al., 2025). Phylogenetic models that support multimodal continuous distributions (Halliwell et al., 2025) will aid understanding how sociosexual selection shapes the adaptive landscapes of social traits across sexes and species.

