## Supplementary Material for "Not all sexual monomorphisms evolve equally: female and male ornamentation in Tyranni passerines"

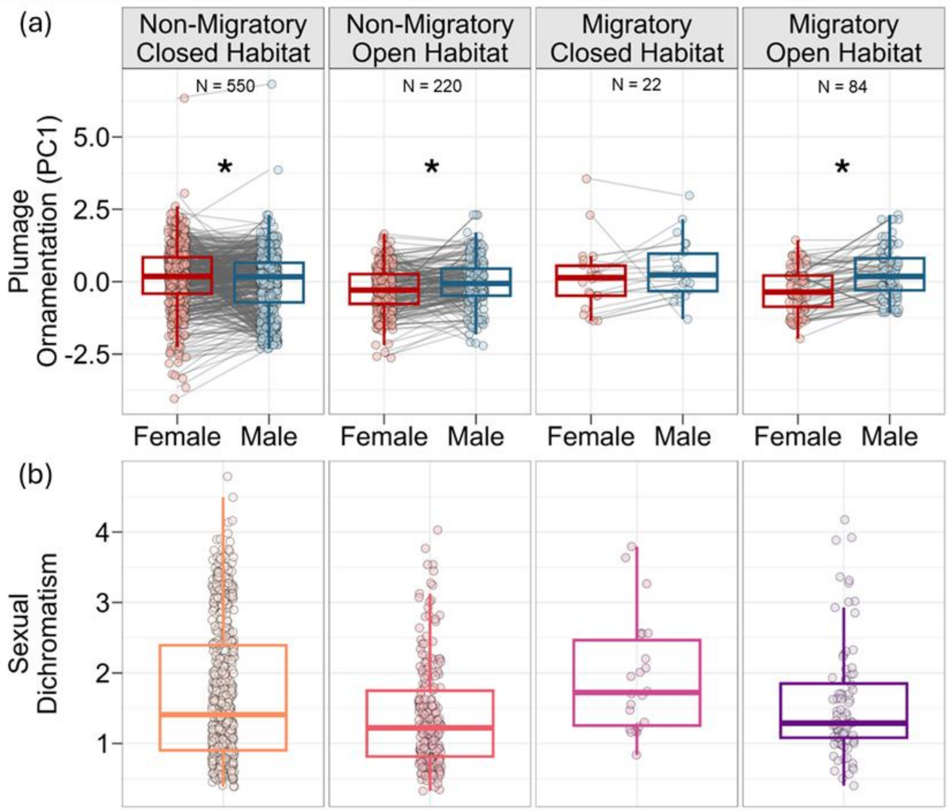


**Figure S1.** Migratory behavior and open habitat types are associated with more dissimilar ornamentation between the sexes. Boxplots are in the style of Tukey and show (a) PC1 of females and males and (b) sexual dichromatism of the interaction between migratory behavior and habitat type. PC1 was obtained with a phylogenetic PCA and represents an axis of increasing plumage color diversity, contrast and saturation in females and males. Sexual dichromatism was calculated with 3D volumes and represents how different females and males are in the avian tetracolor space. Asterisks indicate comparisons that are credibly different from zero. Lines connect points referring to female and male plumage ornamentation of the same species. Numbers in upper plots are sample sizes in each category.

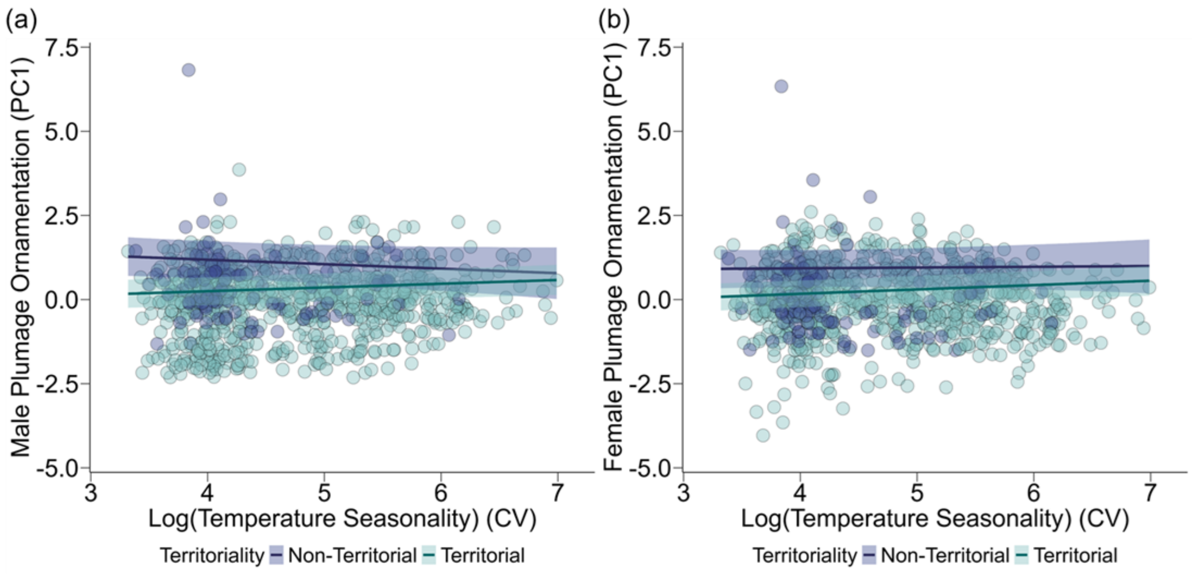


**Figure S2.** Higher climatic seasonality in territorial species is associated with higher (a) male ornamentation, whereas female ornamentation (b) is not credibly associated with temperature seasonality. PC1 was obtained with a phylogenetic PCA and represents an axis of increasing plumage color diversity, contrast and saturation in females and males. Each point represents a species, which are colored according to territoriality categories. Shaded areas are 95% CI.

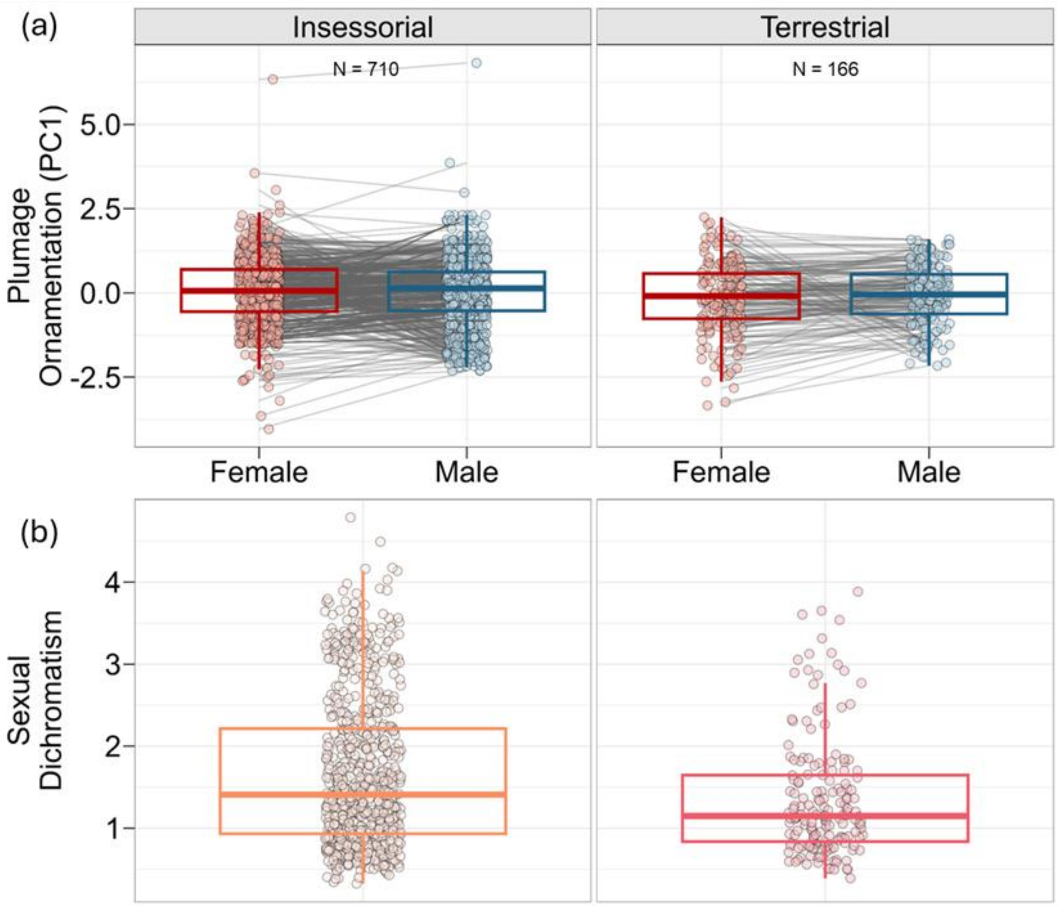


**Figure S3.** Females and males show similar ornamentation in insessorial and terrestrial species. Boxplots are in the style of Tukey and show (a) PC1 of females and males and (b) sexual dichromatism of the locomotion style categories. PC1 was obtained with a phylogenetic PCA and represents an axis of increasing plumage color diversity, contrast and saturation in females and males. Sexual dichromatism was calculated with 3D volumes and represents how different females and males are in the avian tetracolor space. Lines connect points referring to female and male plumage ornamentation of the same species. Numbers in upper plots are sample sizes in each category.

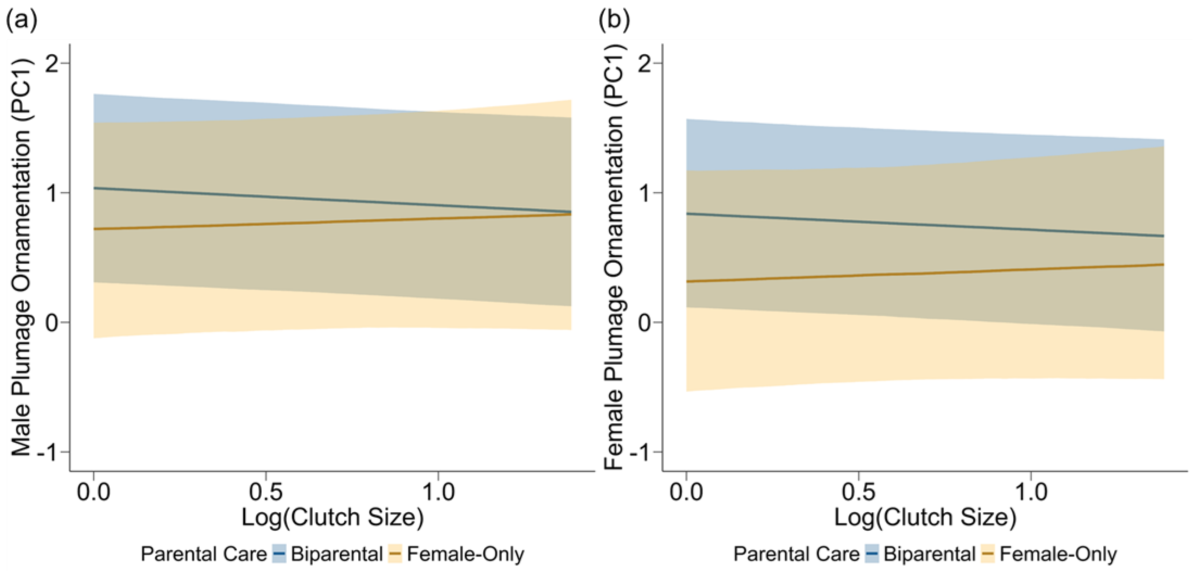


**Figure S4.** As clutch size increases, males (a) show higher ornamentation than females (b) in species with female-only parental care, whereas clutch size does not credibly affect ornamentation in either sex in species with biparental care. PC1 was obtained with a phylogenetic PCA and represents an axis of increasing plumage color diversity, contrast and saturation in females and males. Each point represents a species, which are colored according to territoriality and social mating system categories. Shaded areas are 95% CI.

### Comparisons between the models with and without imputation

The models without imputation of missing data in the predictors showed generally similar results to the models with imputation. However, some effects differed due to changes in sample sizes and credible intervals in addition to possible artifacts introduced by imputation. Namely, the effect of interaction between social mating system and sexual size dimorphism on plumage sexual dichromatism was present when considering PC1 ($\beta$ = 0.89 [0.36 : 1.42]) in the model without imputation, but absent in the model with imputation ($\beta$ = 0.38 [-0.14 : 0.90]; Tables S1 and S6). In post-hoc contrasts, species with long term social pair bonds (PC1: $\beta$ = 0.08 [-0.03 : 0.19]; PC2: $\beta$ = 0.03 [-0.13 : 0.07]), non-migratory territorial species (PC1: $\beta$ = 0.07 [-0.02 : 0.19]; PC2: $\beta$ = 0 [-0.11 : 0.10]), non-migratory species living in closed habitats (PC1: $\beta$ = 0.10 [-0.04 : 0.24]; PC2: $\beta$ = 0.03 [-0.10 : 0.16]), and non-territorial monogamous species (PC1: $\beta = -0.02$ [-0.70 : 0.65]) no longer showed credible effects of higher ornamentation in females than in males; instead females and males showed similar ornamentation in these cases (Tables S9 and S10).

**Table S1.** Results of the Bayesian phylogenetic multilevel model with sexual dichromatism as the response variable.

| Parameters | Estimate | SE | L-95% | U-95% | $\hat{R}$ | Bulk ESS | Tail ESS | $R^2$ |
|---|---|---|---|---|---|---|---|---|
| Intercept | 1.16 | 0.99 | -0.78 | 3.09 | 1.00 | 5909 | 10330 | 0.82 |
| Social Pair Bond: Long Term - Short Term | **-0.10** | **0.04** | **-0.19** | **-0.02** | 1.00 | 8800 | 13451 | |
| Social Pair Bond: Long Term - Solitary | 0.15 | 0.21 | -0.26 | 0.56 | 1.00 | 5437 | 9459 | |
| Habitat Type: Closed - Open | -0.01 | 0.03 | -0.07 | 0.06 | 1.00 | 8638 | 13106 | |
| Migratory Behavior: Absent - Present | **0.43** | **0.14** | **0.15** | **0.71** | 1.00 | 7416 | 11228 | |
| Territoriality: Non-Territorial - Territorial | -0.38 | 0.88 | -2.12 | 1.33 | 1.00 | 4696 | 8035 | |
| Locomotion Style: Insessorial - Terrestrial | 0.03 | 0.05 | -0.06 | 0.13 | 1.00 | 7804 | 11806 | |
| Nest Type: Dome - Primary Cavity | 0.00 | 0.09 | -0.18 | 0.19 | 1.00 | 6955 | 11124 | |
| Nest Type: Dome - Secondary Cavity | 0.02 | 0.07 | -0.10 | 0.15 | 1.00 | 5492 | 9435 | |
| Nest Type: Dome - Open Cup | -0.09 | 0.07 | -0.22 | 0.04 | 1.00 | 5418 | 9878 | |
| Parental Care: Biparental - Female-Only | **-0.36** | **0.16** | **-0.66** | **-0.05** | 1.00 | 5895 | 10002 | |
| Clutch Size | -0.03 | 0.05 | -0.13 | 0.07 | 1.00 | 9851 | 13555 | |
| Social Mating System: Monogamous - Polygynous | **0.33** | **0.16** | **0.01** | **0.65** | 1.00 | 6525 | 10719 | |
| Wing Length | -0.15 | 0.18 | -0.51 | 0.20 | 1.00 | 4836 | 8310 | |
| Sexual Size Dimorphism | -0.01 | 0.01 | -0.03 | 0.01 | 1.00 | 11123 | 14384 | |
| Net Primary Productivity | -0.05 | 0.04 | -0.12 | 0.02 | 1.00 | 10384 | 13975 | |
| Temperature | 0.02 | 0.05 | -0.08 | 0.12 | 1.00 | 9816 | 13823 | |
| Precipitation | -0.01 | 0.03 | -0.06 | 0.04 | 1.00 | 10258 | 14173 | |
| Temperature Seasonality | 0.01 | 0.05 | -0.10 | 0.11 | 1.00 | 6347 | 11268 | |
| Habitat Type: Closed – Open × Migratory Behavior: Absent - Present | **-0.25** | **0.08** | **-0.41** | **-0.09** | 1.00 | 9445 | 13264 | |
| Migratory Behavior: Absent – Present × Territoriality: Non-Territorial - Territorial | -0.10 | 0.15 | -0.40 | 0.19 | 1.00 | 6555 | 10200 | |
| Nest Type: Dome - Primary Cavity × Parental Care: Biparental - Female-Only | 0.01 | 2.51 | -4.91 | 4.91 | 1.00 | 32120 | 14965 | |
| Nest Type: Dome - Secondary Cavity × Parental Care: Biparental - Female-Only | 0.26 | 0.18 | -0.10 | 0.61 | 1.00 | 6465 | 10024 | |
| Nest Type: Dome - Open Cup × Parental Care: Biparental - Female-Only | **0.37** | **0.17** | **0.04** | **0.71** | 1.00 | 5843 | 10376 | |
| Parental Care: Biparental - Female-Only × Clutch Size | **0.27** | **0.13** | **0.01** | **0.52** | 1.00 | 9522 | 12615 | |
| Territoriality: Non-Territorial – Territorial × Social Mating System: Monogamous - Polygynous | -0.28 | 0.17 | -0.61 | 0.06 | 1.00 | 6660 | 10886 | |
| Territoriality: Non-Territorial – Territorial × Wing Length | 0.15 | 0.19 | -0.22 | 0.52 | 1.00 | 4650 | 8712 | |
| Social Mating System: Monogamous – Polygynous × Sexual Size Dimorphism | 0.00 | 0.03 | -0.06 | 0.06 | 1.00 | 9547 | 14117 | |
| Territoriality: Non-Territorial – Territorial × Temperature Seasonality | -0.01 | 0.06 | -0.12 | 0.10 | 1.00 | 6366 | 10273 | |

**Legend:** SE is the standard error, L- and U-95% are lower and upper bounds of the 95% credible interval, respectively, $\hat{R}$ is the Gelman-Rubin diagnostic, Bulk and Tail ESS are effective sample sizes, and $R^2$ is the coefficient of determination. The sign × denotes an interaction. Values in bold indicate effects whose credible intervals do not cross zero.

**Table S2.1 (continues).** Results of the Bayesian multi-response phylogenetic multilevel model with female PC1 as the response variable.

| Parameters | Estimate | SE | L-95% | U-95% | $\hat{R}$ | Bulk ESS | Tail ESS | $R^2$ |
|---|---|---|---|---|---|---|---|---|
| Intercept | 0.38 | 1.51 | -2.61 | 3.30 | 1.00 | 8036 | 12172 | 0.61 |
| Social Pair Bond: Long Term - Short Term | -0.13 | 0.07 | -0.28 | 0.01 | 1.00 | 13938 | 14414 | |
| Social Pair Bond: Long Term - Solitary | **0.74** | **0.31** | **0.13** | **1.35** | 1.00 | 7990 | 12000 | |
| Habitat Type: Closed - Open | **-0.28** | **0.06** | **-0.39** | **-0.16** | 1.00 | 11971 | 13803 | |
| Migratory Behavior: Absent - Present | 0.37 | 0.24 | -0.10 | 0.84 | 1.00 | 9268 | 12846 | |
| Territoriality: Non-Territorial - Territorial | -2.05 | 1.26 | -4.50 | 0.46 | 1.00 | 6943 | 10236 | |
| Locomotion Style: Insessorial - Terrestrial | 0.03 | 0.08 | -0.13 | 0.18 | 1.00 | 11101 | 13756 | |
| Nest Type: Dome - Primary Cavity | -0.13 | 0.15 | -0.43 | 0.16 | 1.00 | 9431 | 12353 | |
| Nest Type: Dome - Secondary Cavity | 0.03 | 0.10 | -0.18 | 0.23 | 1.00 | 8318 | 12719 | |
| Nest Type: Dome - Open Cup | -0.13 | 0.10 | -0.33 | 0.06 | 1.00 | 8369 | 11842 | |
| Parental Care: Biparental - Female-Only | **-0.52** | **0.26** | **-1.02** | **-0.01** | 1.00 | 7549 | 11768 | |
| Clutch Size | -0.12 | 0.09 | -0.30 | 0.05 | 1.00 | 12988 | 15196 | |
| Social Mating System: Monogamous - Polygynous | **-0.85** | **0.26** | **-1.36** | **-0.32** | 1.00 | 8069 | 12318 | |
| Wing Length | 0.04 | 0.27 | -0.48 | 0.58 | 1.00 | 6994 | 10493 | |
| Sexual Size Dimorphism | 0.01 | 0.02 | -0.02 | 0.05 | 1.00 | 14088 | 14796 | |
| Net Primary Productivity | -0.03 | 0.06 | -0.16 | 0.09 | 1.00 | 14562 | 15448 | |
| Temperature | -0.06 | 0.09 | -0.23 | 0.11 | 1.00 | 12866 | 15268 | |
| Precipitation | **0.11** | **0.05** | **0.02** | **0.21** | 1.00 | 13744 | 15702 | |
| Temperature Seasonality | -0.02 | 0.10 | -0.20 | 0.17 | 1.00 | 7845 | 11768 | |
| Habitat Type: Closed – Open × Migratory Behavior: Absent - Present | 0.03 | 0.14 | -0.26 | 0.31 | 1.00 | 13052 | 14937 | |
| Migratory Behavior: Absent – Present × Territoriality: Non-Territorial - Territorial | -0.39 | 0.25 | -0.90 | 0.10 | 1.00 | 8439 | 12884 | |
| Nest Type: Dome - Primary Cavity × Parental Care: Biparental - Female-Only | -0.02 | 2.51 | -4.92 | 4.84 | 1.00 | 31993 | 13970 | |
| Nest Type: Dome - Secondary Cavity × Parental Care: Biparental - Female-Only | 0.41 | 0.29 | -0.17 | 0.99 | 1.00 | 8042 | 11948 | |
| Nest Type: Dome - Open Cup × Parental Care: Biparental - Female-Only | 0.05 | 0.26 | -0.46 | 0.56 | 1.00 | 7461 | 11296 | |
| Parental Care: Biparental - Female-Only × Clutch Size | 0.22 | 0.23 | -0.22 | 0.66 | 1.00 | 10669 | 14043 | |
| Territoriality: Non-Territorial – Territorial × Social Mating System: Monogamous - Polygynous | **0.91** | **0.28** | **0.36** | **1.45** | 1.00 | 7981 | 12230 | |
| Territoriality: Non-Territorial – Territorial × Wing Length | 0.20 | 0.27 | -0.34 | 0.73 | 1.00 | 6759 | 10285 | |
| Social Mating System: Monogamous – Polygynous × Sexual Size Dimorphism | 0.08 | 0.06 | -0.03 | 0.19 | 1.00 | 12586 | 15592 | |
| Territoriality: Non-Territorial – Territorial × Temperature Seasonality | 0.14 | 0.10 | -0.05 | 0.33 | 1.00 | 8067 | 11605 | |

**Legend:** SE is the standard error, L- and U-95% are lower and upper bounds of the 95% credible interval, respectively, $\hat{R}$ is the Gelman-Rubin diagnostic, Bulk and Tail ESS are effective sample sizes, and $R^2$ is the coefficient of determination. The sign × denotes an interaction. Values in bold indicate effects whose credible intervals do not cross zero.

**Table S2.2 (continuation).** Results of the Bayesian multi-response phylogenetic multilevel model with male PC1 as the response variable.

| Parameters | Estimate | SE | L-95% | U-95% | $\widehat{R}$ | Bulk ESS | Tail ESS | $R^2$ |
|---|---|---|---|---|---|---|---|---|
| Intercept | 0.32 | 1.45 | -2.51 | 3.16 | 1.00 | 6942 | 11122 | 0.72 |
| Social Pair Bond: Long Term - Short Term | **-0.14** | **0.07** | **-0.27** | **-0.01** | 1.00 | 11750 | 14822 | |
| Social Pair Bond: Long Term - Solitary | 0.19 | 0.30 | -0.40 | 0.79 | 1.00 | 7490 | 10888 | |
| Habitat Type: Closed - Open | -0.11 | 0.05 | -0.21 | 0.00 | 1.00 | 11011 | 13598 | |
| Migratory Behavior: Absent - Present | -0.10 | 0.22 | -0.53 | 0.34 | 1.00 | 8461 | 12220 | |
| Territoriality: Non-Territorial - Territorial | -1.84 | 1.24 | -4.25 | 0.58 | 1.00 | 6073 | 9858 | |
| Locomotion Style: Insessorial - Terrestrial | -0.06 | 0.07 | -0.21 | 0.08 | 1.00 | 8910 | 13474 | |
| Nest Type: Dome - Primary Cavity | 0.04 | 0.14 | -0.23 | 0.31 | 1.00 | 8829 | 11907 | |
| Nest Type: Dome - Secondary Cavity | 0.11 | 0.10 | -0.09 | 0.30 | 1.00 | 7189 | 11638 | |
| Nest Type: Dome - Open Cup | -0.10 | 0.10 | -0.29 | 0.09 | 1.00 | 7384 | 11857 | |
| Parental Care: Biparental - Female-Only | -0.32 | 0.24 | -0.81 | 0.14 | 1.00 | 6525 | 10925 | |
| Clutch Size | -0.13 | 0.08 | -0.29 | 0.03 | 1.00 | 12517 | 14649 | |
| Social Mating System: Monogamous - Polygynous | -0.45 | 0.25 | -0.95 | 0.04 | 1.00 | 8137 | 10941 | |
| Wing Length | -0.13 | 0.26 | -0.63 | 0.37 | 1.00 | 5974 | 10069 | |
| Sexual Size Dimorphism | 0.02 | 0.02 | -0.01 | 0.05 | 1.00 | 12494 | 15075 | |
| Net Primary Productivity | 0.11 | 0.06 | -0.01 | 0.22 | 1.00 | 12538 | 14780 | |
| Temperature | 0.11 | 0.08 | -0.05 | 0.26 | 1.00 | 12164 | 14485 | |
| Precipitation | **0.10** | **0.04** | **0.01** | **0.18** | 1.00 | 11958 | 14618 | |
| Temperature Seasonality | -0.15 | 0.09 | -0.32 | 0.02 | 1.00 | 8650 | 12786 | |
| Habitat Type: Closed – Open × Migratory Behavior: Absent - Present | 0.04 | 0.13 | -0.23 | 0.30 | 1.00 | 11747 | 14794 | |
| Migratory Behavior: Absent – Present × Territoriality: Non-Territorial - Territorial | 0.21 | 0.24 | -0.25 | 0.69 | 1.00 | 7750 | 11870 | |
| Nest Type: Dome - Primary Cavity × Parental Care: Biparental - Female-Only | 0.00 | 2.50 | -4.89 | 4.83 | 1.00 | 32471 | 14781 | |
| Nest Type: Dome - Secondary Cavity × Parental Care: Biparental - Female-Only | 0.21 | 0.27 | -0.33 | 0.75 | 1.00 | 7286 | 11662 | |
| Nest Type: Dome - Open Cup × Parental Care: Biparental - Female-Only | 0.01 | 0.25 | -0.49 | 0.51 | 1.00 | 7175 | 11213 | |
| Parental Care: Biparental - Female-Only × Clutch Size | 0.22 | 0.21 | -0.19 | 0.62 | 1.00 | 10623 | 13559 | |
| Territoriality: Non-Territorial – Territorial × Social Mating System: Monogamous - Polygynous | 0.38 | 0.26 | -0.14 | 0.90 | 1.00 | 8180 | 11879 | |
| Territoriality: Non-Territorial – Territorial × Wing Length | 0.00 | 0.27 | -0.52 | 0.52 | 1.00 | 5864 | 10694 | |
| Social Mating System: Monogamous – Polygynous × Sexual Size Dimorphism | 0.09 | 0.05 | -0.01 | 0.19 | 1.00 | 11208 | 14804 | |
| Territoriality: Non-Territorial – Territorial × Temperature Seasonality | **0.26** | **0.09** | **0.09** | **0.44** | 1.00 | 8432 | 12321 | |
| Residual Correlation: Female PC1 – Male PC1 | **0.50** | **0.03** | **0.45** | **0.55** | 1.00 | 5809 | 9735 | |

**Legend:** SE is the standard error, L- and U-95% are lower and upper bounds of the 95% credible interval, respectively, $\widehat{R}$ is the Gelman-Rubin diagnostic, Bulk and Tail ESS are effective sample sizes, and $R^2$ is the coefficient of determination. The sign × denotes an interaction. Values in bold indicate effects whose credible intervals do not cross zero.

**Table S3.1 (continues).** Results of the Bayesian multi-response phylogenetic multilevel model with female PC2 as the response variable.

| Parameters | Estimate | SE | L-95% | U-95% | $\hat{R}$ | Bulk ESS | Tail ESS | $R^2$ |
|---|---|---|---|---|---|---|---|---|
| Intercept | -1.23 | 1.62 | -4.38 | 1.94 | 1.00 | 3582 | 6403 | 0.22 |
| Social Pair Bond: Long Term - Short Term | -0.09 | 0.08 | -0.25 | 0.07 | 1.00 | 7198 | 7362 | |
| Social Pair Bond: Long Term - Solitary | 0.76 | 0.31 | 0.16 | 1.36 | 1.00 | 4269 | 5824 | |
| Habitat Type: Closed - Open | **-0.29** | **0.07** | **-0.41** | **-0.16** | 1.00 | 5944 | 7060 | |
| Migratory Behavior: Absent - Present | **-0.60** | **0.28** | **-1.16** | **-0.05** | 1.00 | 3821 | 6584 | |
| Territoriality: Non-Territorial - Territorial | -0.46 | 1.26 | -2.92 | 2.06 | 1.00 | 2538 | 4476 | |
| Locomotion Style: Insessorial - Terrestrial | 0.14 | 0.08 | -0.02 | 0.30 | 1.00 | 6054 | 6835 | |
| Nest Type: Dome - Primary Cavity | **-0.38** | **0.14** | **-0.65** | **-0.10** | 1.00 | 4335 | 6938 | |
| Nest Type: Dome - Secondary Cavity | **-0.24** | **0.10** | **-0.44** | **-0.05** | 1.00 | 3877 | 6389 | |
| Nest Type: Dome - Open Cup | -0.09 | 0.10 | -0.27 | 0.10 | 1.00 | 4207 | 5658 | |
| Parental Care: Biparental - Female-Only | -0.47 | 0.28 | -1.02 | 0.07 | 1.00 | 3612 | 5967 | |
| Clutch Size | -0.08 | 0.11 | -0.29 | 0.14 | 1.00 | 5796 | 7524 | |
| Social Mating System: Monogamous - Polygynous | -0.32 | 0.29 | -0.89 | 0.27 | 1.00 | 3492 | 5243 | |
| Wing Length | 0.40 | 0.26 | -0.11 | 0.92 | 1.00 | 2692 | 4731 | |
| Sexual Size Dimorphism | -0.02 | 0.02 | -0.06 | 0.03 | 1.00 | 7019 | 7391 | |
| Net Primary Productivity | -0.14 | 0.08 | -0.30 | 0.01 | 1.00 | 6761 | 7681 | |
| Temperature | 0.02 | 0.10 | -0.18 | 0.21 | 1.00 | 6471 | 7235 | |
| Precipitation | -0.01 | 0.06 | -0.12 | 0.11 | 1.00 | 6359 | 7156 | |
| Temperature Seasonality | 0.21 | 0.12 | -0.03 | 0.43 | 1.00 | 3503 | 5378 | |
| Habitat Type: Closed – Open × Migratory Behavior: Absent - Present | 0.35 | 0.18 | 0.00 | 0.70 | 1.00 | 5991 | 6496 | |
| Migratory Behavior: Absent – Present × Territoriality: Non-Territorial - Territorial | 0.28 | 0.30 | -0.32 | 0.85 | 1.00 | 3790 | 5403 | |
| Nest Type: Dome - Primary Cavity × Parental Care: Biparental - Female-Only | -0.03 | 2.54 | -5.05 | 4.85 | 1.00 | 10837 | 7200 | |
| Nest Type: Dome - Secondary Cavity × Parental Care: Biparental - Female-Only | 0.63 | 0.31 | 0.03 | 1.24 | 1.00 | 4095 | 6094 | |
| Nest Type: Dome - Open Cup × Parental Care: Biparental - Female-Only | -0.23 | 0.26 | -0.74 | 0.27 | 1.00 | 3719 | 5850 | |
| Parental Care: Biparental - Female-Only × Clutch Size | 0.46 | 0.26 | -0.05 | 0.98 | 1.00 | 4722 | 6711 | |
| Territoriality: Non-Territorial – Territorial × Social Mating System: Monogamous - Polygynous | 0.20 | 0.30 | -0.39 | 0.79 | 1.00 | 3592 | 5679 | |
| Territoriality: Non-Territorial – Territorial × Wing Length | 0.20 | 0.27 | -0.33 | 0.73 | 1.00 | 2683 | 4494 | |
| Social Mating System: Monogamous – Polygynous × Sexual Size Dimorphism | 0.11 | 0.07 | -0.02 | 0.24 | 1.00 | 6512 | 7133 | |
| Territoriality: Non-Territorial – Territorial × Temperature Seasonality | -0.06 | 0.12 | -0.29 | 0.18 | 1.00 | 3403 | 5381 | |

**Legend:** SE is the standard error, L- and U-95% are lower and upper bounds of the 95% credible interval, respectively, $\hat{R}$ is the Gelman-Rubin diagnostic, Bulk and Tail ESS are effective sample sizes, and $R^2$ is the coefficient of determination. The sign × denotes an interaction. Values in bold indicate effects whose credible intervals do not cross zero.

**Table S3.2 (continuation).** Results of the Bayesian multi-response phylogenetic multilevel model with male PC2 as the response variable.

| Parameters | Estimate | SE | L-95% | U-95% | $\hat{R}$ | Bulk ESS | Tail ESS | $R^2$ |
|---|---|---|---|---|---|---|---|---|
| Intercept | -1.24 | 1.45 | -4.07 | 1.59 | 1.00 | 3580 | 5748 | 0.44 |
| Social Pair Bond: Long Term - Short Term | -0.13 | 0.07 | -0.27 | 0.02 | 1.00 | 6776 | 7319 | |
| Social Pair Bond: Long Term - Solitary | 0.40 | 0.28 | -0.14 | 0.94 | 1.00 | 3860 | 5729 | |
| Habitat Type: Closed - Open | **-0.17** | **0.06** | **-0.28** | **-0.05** | 1.00 | 5628 | 6881 | |
| Migratory Behavior: Absent - Present | **-0.74** | **0.25** | **-1.23** | **-0.25** | 1.00 | 3965 | 5961 | |
| Territoriality: Non-Territorial - Territorial | -0.30 | 1.17 | -2.58 | 2.03 | 1.00 | 2521 | 4483 | |
| Locomotion Style: Insessorial - Terrestrial | 0.02 | 0.07 | -0.12 | 0.17 | 1.00 | 6006 | 7490 | |
| Nest Type: Dome - Primary Cavity | **-0.30** | **0.13** | **-0.55** | **-0.04** | 1.00 | 4377 | 6420 | |
| Nest Type: Dome - Secondary Cavity | -0.08 | 0.09 | -0.26 | 0.09 | 1.00 | 4141 | 6418 | |
| Nest Type: Dome - Open Cup | -0.03 | 0.09 | -0.20 | 0.14 | 1.00 | 4276 | 6205 | |
| Parental Care: Biparental - Female-Only | -0.38 | 0.25 | -0.87 | 0.10 | 1.00 | 3072 | 4939 | |
| Clutch Size | -0.15 | 0.10 | -0.34 | 0.04 | 1.00 | 6865 | 7353 | |
| Social Mating System: Monogamous - Polygynous | -0.13 | 0.26 | -0.64 | 0.37 | 1.00 | 3364 | 5081 | |
| Wing Length | 0.11 | 0.24 | -0.36 | 0.59 | 1.00 | 2660 | 4338 | |
| Sexual Size Dimorphism | -0.01 | 0.02 | -0.05 | 0.03 | 1.00 | 6810 | 7849 | |
| Net Primary Productivity | -0.02 | 0.07 | -0.16 | 0.12 | 1.00 | 6571 | 7689 | |
| Temperature | 0.14 | 0.09 | -0.04 | 0.31 | 1.00 | 6571 | 7279 | |
| Precipitation | 0.03 | 0.05 | -0.07 | 0.13 | 1.00 | 6286 | 7235 | |
| Temperature Seasonality | 0.15 | 0.10 | -0.06 | 0.34 | 1.00 | 3452 | 5427 | |
| Habitat Type: Closed – Open × Migratory Behavior: Absent - Present | 0.22 | 0.15 | -0.08 | 0.52 | 1.00 | 5518 | 6387 | |
| Migratory Behavior: Absent – Present × Territoriality: Non-Territorial - Territorial | 0.69 | 0.27 | 0.17 | 1.21 | 1.00 | 3482 | 4999 | |
| Nest Type: Dome - Primary Cavity × Parental Care: Biparental - Female-Only | 0.01 | 2.49 | -4.89 | 4.90 | 1.00 | 10772 | 7534 | |
| Nest Type: Dome - Secondary Cavity × Parental Care: Biparental - Female-Only | **0.62** | **0.27** | **0.09** | **1.16** | 1.00 | 3852 | 6116 | |
| Nest Type: Dome - Open Cup × Parental Care: Biparental - Female-Only | 0.07 | 0.23 | -0.39 | 0.52 | 1.00 | 3723 | 6218 | |
| Parental Care: Biparental - Female-Only × Clutch Size | 0.23 | 0.23 | -0.23 | 0.69 | 1.00 | 4253 | 6362 | |
| Territoriality: Non-Territorial – Territorial × Social Mating System: Monogamous - Polygynous | -0.15 | 0.27 | -0.68 | 0.37 | 1.00 | 3545 | 5506 | |
| Territoriality: Non-Territorial – Territorial × Wing Length | 0.09 | 0.25 | -0.40 | 0.58 | 1.00 | 2566 | 4023 | |
| Social Mating System: Monogamous – Polygynous × Sexual Size Dimorphism | **0.15** | **0.06** | **0.03** | **0.27** | 1.00 | 6027 | 6481 | |
| Territoriality: Non-Territorial – Territorial × Temperature Seasonality | -0.03 | 0.11 | -0.23 | 0.18 | 1.00 | 3381 | 5589 | |
| Residual Correlation: Female PC2 – Male PC2 | **0.37** | **0.02** | **0.33** | **0.41** | 1.00 | 7185 | 7371 | |

**Legend:** SE is the standard error, L- and U-95% are lower and upper bounds of the 95% credible interval, respectively, $\hat{R}$ is the Gelman-Rubin diagnostic, Bulk and Tail ESS are effective sample sizes, and $R^2$ is the coefficient of determination. The sign × denotes an interaction. Values in bold indicate effects whose credible intervals do not cross zero.

**Table S4.** Contrasts between female and male PC1 in each parameter category.

| Parameters | Contrast (Female - Male) | Estimate | L-95% | U-95% |
|---|---|---|---|---|
| Social Pair Bond | Long-Term | **0.16** | **0.07** | **0.24** |
| | Short-Term | **-0.43** | **-0.62** | **-0.24** |
| | Solitary | **-0.56** | **-0.85** | **-0.27** |
| Territoriality × Social Mating System | Non-Territorial : Monogamous | **0.58** | **0.04** | **1.11** |
| | Territorial : Monogamous | 0.06 | -0.02 | 0.14 |
| | Non-Territorial : Polygynous | **-0.58** | **-0.86** | **-0.30** |
| | Territorial : Polygynous | -0.34 | -0.74 | 0.07 |
| Territoriality × Migratory Behavior | Non-Territorial : Migratory | 0.14 | -0.57 | 0.89 |
| | Territorial : Migratory | **-0.49** | **-0.73** | **-0.26** |
| | Non-Territorial : Non-Migratory | **-0.41** | **-0.70** | **-0.13** |
| | Territorial : Non-Migratory | **0.13** | **0.04** | **0.20** |
| Parental Care × Nest Type | Biparental : Primary Cavity | -0.24 | -0.54 | 0.04 |
| | Biparental : Secondary Cavity | **-0.24** | **-0.41** | **-0.07** |
| | Female-Only : Secondary Cavity | -0.13 | -0.75 | 0.48 |
| | Biparental : Dome | -0.10 | -0.25 | 0.06 |
| | Female-Only : Dome | **-0.61** | **-1.17** | **-0.06** |
| | Biparental × Open Cup | **0.36** | **0.25** | **0.47** |
| | Female-Only : Open Cup | **-0.59** | **-0.87** | **-0.32** |
| Migratory Behavior × Habitat Type | Non-Migratory : Closed Habitat | **0.17** | **0.07** | **0.27** |
| | Migratory : Closed Habitat | -0.26 | -0.71 | 0.19 |
| | Non-Migratory : Open Habitat | **-0.20** | **-0.36** | **-0.05** |
| | Migratory : Open Habitat | **-0.50** | **-0.74** | **-0.24** |
| Locomotion Style | Insessorial | -0.01 | -0.18 | 0.16 |
| | Terrestrial | 0.00 | -0.08 | 0.09 |

**Legend:** L- and U-95% are lower and upper bounds of the 95% credible interval, respectively. The sign × denotes an interaction. Values in bold indicate effects whose credible intervals do not cross zero.

**Table S5.** Contrasts between female and male PC2 in each parameter category.

| Parameters | Contrast (Female - Male) | Estimate | L-95% | U-95% |
|---|---|---|---|---|
| Social Pair Bond | Long-Term | **0.09** | **0.01** | **0.17** |
| | Short-Term | **-0.21** | **-0.39** | **-0.02** |
| | Solitary | **-0.37** | **-0.65** | **-0.10** |
| Territoriality × Social Mating System | Non-Territorial : Monogamous | 0.10 | -0.41 | 0.61 |
| | Territorial : Monogamous | 0.04 | -0.03 | 0.12 |
| | Non-Territorial : Polygynous | **-0.39** | **-0.66** | **-0.12** |
| | Territorial : Polygynous | -0.03 | -0.42 | 0.36 |
| Territoriality × Migratory Behavior | Non-Territorial : Migratory | -0.14 | -0.83 | 0.57 |
| | Territorial : Migratory | **-0.35** | **-0.58** | **-0.13** |
| | Non-Territorial : Non-Migratory | **-0.31** | **-0.58** | **-0.04** |
| | Territorial : Non-Migratory | **0.10** | **0.02** | **0.18** |
| Parental Care × Nest Type | Biparental : Primary Cavity | -0.07 | -0.35 | 0.20 |
| | Biparental : Secondary Cavity | -0.16 | -0.33 | 0.00 |
| | Female-Only : Secondary Cavity | 0.12 | -0.47 | 0.70 |
| | Biparental : Dome | -0.13 | -0.28 | 0.02 |
| | Female-Only : Dome | -0.22 | -0.73 | 0.30 |
| | Biparental × Open Cup | **0.25** | **0.14** | **0.36** |
| | Female-Only : Open Cup | **-0.40** | **-0.66** | **-0.13** |
| Migratory Behavior × Habitat Type | Non-Migratory : Closed Habitat | **0.14** | **0.04** | **0.23** |
| | Migratory : Closed Habitat | -0.33 | -0.76 | 0.12 |
| | Non-Migratory : Open Habitat | **-0.17** | **-0.32** | **-0.02** |
| | Migratory : Open Habitat | **-0.34** | **-0.57** | **-0.09** |
| Locomotion Style | Insessorial | 0.02 | -0.15 | 0.19 |
| | Terrestrial | 0.00 | -0.08 | 0.08 |

**Legend:** L- and U-95% are lower and upper bounds of the 95% credible interval, respectively. The sign × denotes an interaction. Values in bold indicate effects whose credible intervals do not cross zero.

**Table S6.** Results of the Bayesian phylogenetic multilevel model with sexual dichromatism as the response variable and without imputation of missing data in the predictors.

| Parameters | Estimate | SE | L-95% | U-95% | $\hat{R}$ | Bulk ESS | Tail ESS | $R^2$ |
|---|---|---|---|---|---|---|---|---|
| Intercept | 1.03 | 1.11 | -1.17 | 3.20 | 1.00 | 4570 | 8243 | 0.90 |
| Social Pair Bond: Long Term - Short Term | **-0.15** | **0.05** | **-0.24** | **-0.05** | 1.00 | 5673 | 10358 | |
| Social Pair Bond: Long Term - Solitary | 0.13 | 0.22 | -0.30 | 0.56 | 1.00 | 4814 | 8037 | |
| Habitat Type: Closed - Open | 0.03 | 0.04 | -0.05 | 0.12 | 1.00 | 5182 | 9125 | |
| Migratory Behavior: Absent - Present | 0.27 | 0.14 | -0.01 | 0.55 | 1.00 | 6105 | 9551 | |
| Territoriality: Non-Territorial - Territorial | 1.18 | 0.95 | -0.70 | 3.06 | 1.00 | 3919 | 7243 | |
| Locomotion Style: Insessorial - Terrestrial | 0.08 | 0.07 | -0.05 | 0.21 | 1.00 | 4768 | 8300 | |
| Nest Type: Dome - Primary Cavity | 0.07 | 0.11 | -0.14 | 0.28 | 1.00 | 4816 | 8590 | |
| Nest Type: Dome - Secondary Cavity | 0.06 | 0.09 | -0.11 | 0.22 | 1.00 | 4013 | 7785 | |
| Nest Type: Dome - Open Cup | -0.05 | 0.08 | -0.20 | 0.09 | 1.00 | 3728 | 7626 | |
| Parental Care: Biparental - Female-Only | -0.31 | 0.16 | -0.62 | 0.00 | 1.00 | 4476 | 8679 | |
| Clutch Size | 0.02 | 0.05 | -0.09 | 0.13 | 1.00 | 5887 | 10035 | |
| Social Mating System: Monogamous - Polygynous | 0.22 | 0.20 | -0.17 | 0.60 | 1.00 | 4176 | 8327 | |
| Wing Length | 0.00 | 0.20 | -0.38 | 0.40 | 1.00 | 3899 | 7423 | |
| Sexual Size Dimorphism | -0.01 | 0.01 | -0.04 | 0.02 | 1.00 | 6304 | 9423 | |
| Net Primary Productivity | **-0.11** | **0.04** | **-0.20** | **-0.03** | 1.00 | 6686 | 10536 | |
| Temperature | 0.03 | 0.06 | -0.09 | 0.14 | 1.00 | 6492 | 10724 | |
| Precipitation | -0.05 | 0.04 | -0.12 | 0.02 | 1.00 | 6074 | 10408 | |
| Temperature Seasonality | 0.06 | 0.06 | -0.07 | 0.18 | 1.00 | 5216 | 9120 | |
| Habitat Type: Closed – Open x Migratory Behavior: Absent - Present | **-0.35** | **0.08** | **-0.51** | **-0.19** | 1.00 | 6182 | 10329 | |
| Migratory Behavior: Absent – Present x Territoriality: Non-Territorial - Territorial | 0.07 | 0.16 | -0.23 | 0.38 | 1.00 | 5738 | 9080 | |
| Nest Type: Dome - Primary Cavity x Parental Care: Biparental - Female-Only | 0.01 | 2.52 | -4.94 | 4.91 | 1.00 | 27447 | 15906 | |
| Nest Type: Dome - Secondary Cavity x Parental Care: Biparental - Female-Only | 0.22 | 0.18 | -0.14 | 0.57 | 1.00 | 4935 | 8578 | |
| Nest Type: Dome - Open Cup x Parental Care: Biparental - Female-Only | **0.40** | **0.18** | **0.05** | **0.75** | 1.00 | 4560 | 8243 | |
| Parental Care: Biparental - Female-Only x Clutch Size | 0.25 | 0.14 | -0.02 | 0.53 | 1.00 | 6394 | 10196 | |
| Territoriality: Non-Territorial – Territorial x Social Mating System: Monogamous - Polygynous | -0.16 | 0.20 | -0.56 | 0.23 | 1.00 | 4350 | 8067 | |
| Territoriality: Non-Territorial – Territorial x Wing Length | -0.14 | 0.20 | -0.54 | 0.26 | 1.00 | 3688 | 7613 | |
| Social Mating System: Monogamous – Polygynous x Sexual Size Dimorphism | -0.06 | 0.03 | -0.13 | 0 | 1.00 | 6761 | 11426 | |
| Territoriality: Non-Territorial – Territorial x Temperature Seasonality | -0.08 | 0.07 | -0.20 | 0.05 | 1.00 | 5129 | 8376 | |

**Legend:** L- and U-95% are lower and upper bounds of the 95% credible interval, respectively. The sign × denotes an interaction. Values in bold indicate effects whose credible intervals do not cross zero.

**Table S7.1 (continues).** Results of the Bayesian multi-response phylogenetic multilevel model with female PC1 as the response variable and without imputation of missing data in the predictors.

| Parameters | Estimate | SE | L-95% | U-95% | $\hat{R}$ | Bulk ESS | Tail ESS | $R^2$ |
|---|---|---|---|---|---|---|---|---|
| Intercept | 1.26 | 1.51 | -1.73 | 4.20 | 1.00 | 10490 | 14261 | 0.57 |
| Social Pair Bond: Long Term - Short Term | **-0.15** | **0.07** | **-0.30** | **-0.01** | 1.00 | 16311 | 15932 | |
| Social Pair Bond: Long Term - Solitary | 0.48 | 0.28 | -0.07 | 1.03 | 1.00 | 9124 | 12834 | |
| Habitat Type: Closed - Open | **-0.23** | **0.06** | **-0.35** | **-0.11** | 1.00 | 13963 | 15587 | |
| Migratory Behavior: Absent - Present | 0.40 | 0.22 | -0.04 | 0.85 | 1.00 | 10596 | 14033 | |
| Territoriality: Non-Territorial - Territorial | -2.32 | 1.19 | -4.66 | 0.01 | 1.00 | 7817 | 11868 | |
| Locomotion Style: Insessorial - Terrestrial | -0.04 | 0.09 | -0.22 | 0.14 | 1.00 | 13030 | 14871 | |
| Nest Type: Dome - Primary Cavity | **-0.32** | **0.15** | **-0.61** | **-0.03** | 1.00 | 12125 | 13631 | |
| Nest Type: Dome - Secondary Cavity | -0.13 | 0.11 | -0.34 | 0.10 | 1.00 | 10469 | 13155 | |
| Nest Type: Dome - Open Cup | **-0.29** | **0.10** | **-0.47** | **-0.10** | 1.00 | 10754 | 12904 | |
| Parental Care: Biparental - Female-Only | -0.42 | 0.22 | -0.86 | 0.01 | 1.00 | 9518 | 13142 | |
| Clutch Size | -0.07 | 0.09 | -0.24 | 0.10 | 1.00 | 15763 | 16343 | |
| Social Mating System: Monogamous - Polygynous | **-0.70** | **0.27** | **-1.22** | **-0.19** | 1.00 | 8998 | 13143 | |
| Wing Length | 0.15 | 0.25 | -0.33 | 0.63 | 1.00 | 8156 | 11689 | |
| Sexual Size Dimorphism | 0.04 | 0.02 | -0.01 | 0.08 | 1.00 | 14761 | 15788 | |
| Net Primary Productivity | -0.12 | 0.07 | -0.25 | 0.02 | 1.00 | 15541 | 16580 | |
| Temperature | 0.00 | 0.09 | -0.19 | 0.18 | 1.00 | 15728 | 16452 | |
| Precipitation | 0.05 | 0.06 | -0.06 | 0.17 | 1.00 | 15284 | 15474 | |
| Temperature Seasonality | -0.05 | 0.10 | -0.25 | 0.14 | 1.00 | 10018 | 13219 | |
| Habitat Type: Closed – Open x Migratory Behavior: Absent - Present | -0.10 | 0.13 | -0.36 | 0.16 | 1.00 | 14443 | 15373 | |
| Migratory Behavior: Absent – Present x Territoriality: Non-Territorial - Territorial | -0.41 | 0.24 | -0.88 | 0.07 | 1.00 | 9849 | 13082 | |
| Nest Type: Dome - Primary Cavity x Parental Care: Biparental - Female-Only | 0.00 | 2.44 | -4.78 | 4.73 | 1.00 | 42592 | 15800 | |
| Nest Type: Dome - Secondary Cavity x Parental Care: Biparental - Female-Only | 0.29 | 0.25 | -0.21 | 0.78 | 1.00 | 10876 | 14974 | |
| Nest Type: Dome - Open Cup x Parental Care: Biparental - Female-Only | 0.18 | 0.23 | -0.27 | 0.65 | 1.00 | 10600 | 13555 | |
| Parental Care: Biparental - Female-Only x Clutch Size | 0.12 | 0.22 | -0.32 | 0.55 | 1.00 | 12082 | 14815 | |
| Territoriality: Non-Territorial – Territorial x Social Mating System: Monogamous - Polygynous | **0.94** | **0.27** | **0.40** | **1.46** | 1.00 | 9053 | 13364 | |
| Territoriality: Non-Territorial – Territorial x Wing Length | 0.27 | 0.25 | -0.23 | 0.76 | 1.00 | 7608 | 10595 | |
| Social Mating System: Monogamous – Polygynous x Sexual Size Dimorphism | -0.04 | 0.05 | -0.15 | 0.07 | 1.00 | 14355 | 15350 | |
| Territoriality: Non-Territorial – Territorial x Temperature Seasonality | 0.14 | 0.10 | -0.06 | 0.34 | 1.00 | 9671 | 13341 | |

**Legend:** SE is the standard error, L- and U-95% are lower and upper bounds of the 95% credible interval, respectively, $\hat{R}$ is the Gelman-Rubin diagnostic, Bulk and Tail ESS are effective sample sizes, and $R^2$ is the coefficient of determination. The sign × denotes an interaction. Values in bold indicate effects whose credible intervals do not cross zero.

**Table S7.2 (continuation).** Results of the Bayesian multi-response phylogenetic multilevel model with male PC1 as the response variable and without imputation of missing data in the predictors.

| Parameters | Estimate | SE | L-95% | U-95% | $\widehat{R}$ | Bulk ESS | Tail ESS | $R^2$ |
|---|---|---|---|---|---|---|---|---|
| Intercept | 1.56 | 1.53 | -1.45 | 4.55 | 1.00 | 9868 | 12948 | 0.64 |
| Social Pair Bond: Long Term - Short Term | **-0.23** | **0.07** | **-0.37** | **-0.09** | 1.00 | 16078 | 15373 | |
| Social Pair Bond: Long Term - Solitary | 0.28 | 0.28 | -0.28 | 0.84 | 1.00 | 9572 | 13388 | |
| Habitat Type: Closed - Open | -0.11 | 0.06 | -0.23 | 0.02 | 1.00 | 14323 | 16074 | |
| Migratory Behavior: Absent - Present | 0.11 | 0.22 | -0.34 | 0.55 | 1.00 | 10211 | 13715 | |
| Territoriality: Non-Territorial - Territorial | **-4.28** | **1.22** | **-6.67** | **-1.88** | 1.00 | 7531 | 11577 | |
| Locomotion Style: Insessorial - Terrestrial | -0.18 | 0.09 | -0.36 | 0.00 | 1.00 | 12925 | 14941 | |
| Nest Type: Dome - Primary Cavity | 0.00 | 0.15 | -0.29 | 0.29 | 1.00 | 11718 | 14900 | |
| Nest Type: Dome - Secondary Cavity | 0.15 | 0.11 | -0.06 | 0.36 | 1.00 | 10528 | 14029 | |
| Nest Type: Dome - Open Cup | -0.06 | 0.10 | -0.24 | 0.14 | 1.00 | 11368 | 13661 | |
| Parental Care: Biparental - Female-Only | -0.09 | 0.22 | -0.53 | 0.35 | 1.00 | 9515 | 12501 | |
| Clutch Size | -0.09 | 0.09 | -0.26 | 0.08 | 1.00 | 15396 | 16017 | |
| Social Mating System: Monogamous - Polygynous | **-0.80** | **0.27** | **-1.34** | **-0.28** | 1.00 | 9025 | 13263 | |
| Wing Length | -0.23 | 0.25 | -0.72 | 0.26 | 1.00 | 7662 | 11417 | |
| Sexual Size Dimorphism | 0.05 | 0.02 | 0.00 | 0.10 | 1.00 | 14361 | 15830 | |
| Net Primary Productivity | **0.15** | **0.07** | **0.01** | **0.29** | 1.00 | 16010 | 16338 | |
| Temperature | **0.25** | **0.09** | **0.08** | **0.43** | 1.00 | 14834 | 15600 | |
| Precipitation | -0.04 | 0.06 | -0.16 | 0.07 | 1.00 | 15421 | 15807 | |
| Temperature Seasonality | **-0.21** | **0.10** | **-0.40** | **-0.01** | 1.00 | 9956 | 13075 | |
| Habitat Type: Closed – Open x Migratory Behavior: Absent - Present | -0.20 | 0.13 | -0.46 | 0.06 | 1.00 | 14048 | 15623 | |
| Migratory Behavior: Absent – Present x Territoriality: Non-Territorial - Territorial | 0.11 | 0.24 | -0.36 | 0.58 | 1.00 | 9302 | 13128 | |
| Nest Type: Dome - Primary Cavity x Parental Care: Biparental - Female-Only | -0.02 | 2.50 | -4.85 | 4.85 | 1.00 | 40858 | 15052 | |
| Nest Type: Dome - Secondary Cavity x Parental Care: Biparental - Female-Only | 0.03 | 0.25 | -0.46 | 0.52 | 1.00 | 10129 | 14429 | |
| Nest Type: Dome - Open Cup x Parental Care: Biparental - Female-Only | 0.00 | 0.23 | -0.46 | 0.46 | 1.00 | 10429 | 13610 | |
| Parental Care: Biparental - Female-Only x Clutch Size | -0.12 | 0.22 | -0.55 | 0.31 | 1.00 | 11253 | 13107 | |
| Territoriality: Non-Territorial – Territorial x Social Mating System: Monogamous - Polygynous | 0.89 | 0.27 | 0.36 | 1.42 | 1.00 | 9174 | 13191 | |
| Territoriality: Non-Territorial – Territorial x Wing Length | 0.42 | 0.26 | -0.08 | 0.93 | 1.00 | 7383 | 10667 | |
| Social Mating System: Monogamous – Polygynous x Sexual Size Dimorphism | -0.02 | 0.05 | -0.13 | 0.09 | 1.00 | 13449 | 15801 | |
| Territoriality: Non-Territorial – Territorial x Temperature Seasonality | **0.32** | **0.10** | **0.12** | **0.52** | 1.00 | 9774 | 13132 | |
| Residual Correlation: Female PC1 x Male PC1 | **0.56** | **0.03** | **0.50** | **0.61** | 1.00 | 5404 | 8622 | |

**Legend:** SE is the standard error, L- and U-95% are lower and upper bounds of the 95% credible interval, respectively, $\widehat{R}$ is the Gelman-Rubin diagnostic, Bulk and Tail ESS are effective sample sizes, and $R^2$ is the coefficient of determination. The sign × denotes an interaction. Values in bold indicate effects whose credible intervals do not cross zero.

**Table S8.1 (continues).** Results of the Bayesian multi-response phylogenetic multilevel model with female PC2 as the response variable and without imputation of missing data in the predictors.

| Parameters | Estimate | SE | L-95% | U-95% | $\hat{R}$ | Bulk ESS | Tail ESS | $R^2$ |
|---|---|---|---|---|---|---|---|---|
| Intercept | -0.96 | 1.66 | -4.20 | 2.29 | 1.00 | 26223 | 17587 | 0.32 |
| Social Pair Bond: Long Term - Short Term | -0.01 | 0.08 | -0.16 | 0.15 | 1.00 | 33374 | 16786 | |
| Social Pair Bond: Long Term - Solitary | **0.67** | **0.29** | **0.10** | **1.25** | 1.00 | 24170 | 17013 | |
| Habitat Type: Closed - Open | -0.13 | 0.07 | -0.27 | 0.01 | 1.00 | 29789 | 17139 | |
| Migratory Behavior: Absent - Present | **-0.68** | **0.26** | **-1.19** | **-0.17** | 1.00 | 25890 | 16285 | |
| Territoriality: Non-Territorial - Territorial | 0.19 | 1.25 | -2.29 | 2.66 | 1.00 | 19330 | 15704 | |
| Locomotion Style: Insessorial - Terrestrial | 0.27 | 0.10 | 0.08 | 0.45 | 1.00 | 30412 | 16330 | |
| Nest Type: Dome - Primary Cavity | -0.38 | 0.16 | -0.69 | -0.07 | 1.00 | 28842 | 15642 | |
| Nest Type: Dome - Secondary Cavity | -0.34 | 0.12 | -0.57 | -0.11 | 1.00 | 25101 | 17111 | |
| Nest Type: Dome - Open Cup | -0.09 | 0.10 | -0.29 | 0.10 | 1.00 | 25853 | 16513 | |
| Parental Care: Biparental - Female-Only | -0.43 | 0.24 | -0.91 | 0.05 | 1.00 | 22243 | 17057 | |
| Clutch Size | -0.16 | 0.10 | -0.35 | 0.03 | 1.00 | 30507 | 17797 | |
| Social Mating System: Monogamous - Polygynous | -0.01 | 0.29 | -0.57 | 0.56 | 1.00 | 22075 | 16340 | |
| Wing Length | 0.31 | 0.25 | -0.19 | 0.80 | 1.00 | 19173 | 16238 | |
| Sexual Size Dimorphism | -0.05 | 0.03 | -0.11 | 0.00 | 1.00 | 32149 | 17028 | |
| Net Primary Productivity | **-0.20** | **0.08** | **-0.35** | **-0.04** | 1.00 | 33511 | 16382 | |
| Temperature | 0.07 | 0.11 | -0.14 | 0.28 | 1.00 | 34625 | 16484 | |
| Precipitation | -0.03 | 0.07 | -0.16 | 0.10 | 1.00 | 33096 | 15594 | |
| Temperature Seasonality | **0.31** | **0.12** | **0.08** | **0.54** | 1.00 | 21448 | 15993 | |
| Habitat Type: Closed – Open x Migratory Behavior: Absent - Present | 0.10 | 0.15 | -0.20 | 0.41 | 1.00 | 30871 | 16746 | |
| Migratory Behavior: Absent – Present x Territoriality: Non-Territorial - Territorial | **0.55** | **0.28** | **0.01** | **1.09** | 1.00 | 23772 | 16463 | |
| Nest Type: Dome - Primary Cavity x Parental Care: Biparental - Female-Only | -0.01 | 2.49 | -4.93 | 4.88 | 1.00 | 49483 | 13023 | |
| Nest Type: Dome - Secondary Cavity x Parental Care: Biparental - Female-Only | **0.53** | **0.27** | **0.01** | **1.06** | 1.00 | 24596 | 16882 | |
| Nest Type: Dome - Open Cup x Parental Care: Biparental - Female-Only | -0.17 | 0.24 | -0.65 | 0.31 | 1.00 | 24957 | 17119 | |
| Parental Care: Biparental - Female-Only x Clutch Size | 0.35 | 0.25 | -0.13 | 0.83 | 1.00 | 25321 | 16383 | |
| Territoriality: Non-Territorial – Territorial x Social Mating System: Monogamous - Polygynous | 0.00 | 0.29 | -0.57 | 0.57 | 1.00 | 22280 | 16414 | |
| Territoriality: Non-Territorial – Territorial x Wing Length | 0.23 | 0.26 | -0.29 | 0.75 | 1.00 | 19240 | 15774 | |
| Social Mating System: Monogamous – Polygynous x Sexual Size Dimorphism | 0.06 | 0.06 | -0.06 | 0.18 | 1.00 | 31521 | 17034 | |
| Territoriality: Non-Territorial – Territorial x Temperature Seasonality | -0.19 | 0.12 | -0.43 | 0.04 | 1.00 | 21326 | 15855 | |

**Legend:** SE is the standard error, L- and U-95% are lower and upper bounds of the 95% credible interval, respectively, $\hat{R}$ is the Gelman-Rubin diagnostic, Bulk and Tail ESS are effective sample sizes, and $R^2$ is the coefficient of determination. The sign × denotes an interaction. Values in bold indicate effects whose credible intervals do not cross zero.

**Table S8.2 (continues).** Results of the Bayesian multi-response phylogenetic multilevel model with male PC2 as the response variable and without imputation of missing data in the predictors.

| Parameters | Estimate | SE | L-95% | U-95% | $\hat{R}$ | Bulk ESS | Tail ESS | $R^2$ |
|---|---|---|---|---|---|---|---|---|
| Intercept | -2.02 | 1.61 | -5.18 | 1.15 | 1.00 | 24291 | 16930 | 0.47 |
| Social Pair Bond: Long Term - Short Term | -0.07 | 0.08 | -0.22 | 0.08 | 1.00 | 34606 | 16464 | |
| Social Pair Bond: Long Term - Solitary | **0.74** | **0.29** | **0.17** | **1.32** | 1.00 | 23122 | 17334 | |
| Habitat Type: Closed - Open | -0.12 | 0.07 | -0.25 | 0.02 | 1.00 | 29207 | 17377 | |
| Migratory Behavior: Absent - Present | **-0.86** | **0.25** | **-1.35** | **-0.37** | 1.00 | 24218 | 16494 | |
| Territoriality: Non-Territorial - Territorial | -0.44 | 1.24 | -2.87 | 1.98 | 1.00 | 17852 | 15848 | |
| Locomotion Style: Insessorial - Terrestrial | 0.06 | 0.09 | -0.12 | 0.24 | 1.00 | 29589 | 16943 | |
| Nest Type: Dome - Primary Cavity | -0.28 | 0.15 | -0.57 | 0.02 | 1.00 | 29377 | 16658 | |
| Nest Type: Dome - Secondary Cavity | -0.21 | 0.11 | -0.43 | 0.01 | 1.00 | 24580 | 16435 | |
| Nest Type: Dome - Open Cup | 0.01 | 0.10 | -0.17 | 0.20 | 1.00 | 26454 | 17795 | |
| Parental Care: Biparental - Female-Only | -0.27 | 0.24 | -0.74 | 0.19 | 1.00 | 22651 | 16843 | |
| Clutch Size | **-0.24** | **0.10** | **-0.43** | **-0.05** | 1.00 | 32134 | 17179 | |
| Social Mating System: Monogamous - Polygynous | 0.07 | 0.29 | -0.49 | 0.64 | 1.00 | 22214 | 17163 | |
| Wing Length | 0.07 | 0.25 | -0.43 | 0.56 | 1.00 | 18317 | 15969 | |
| Sexual Size Dimorphism | -0.05 | 0.03 | -0.10 | 0.01 | 1.00 | 33920 | 18142 | |
| Net Primary Productivity | 0.07 | 0.08 | -0.08 | 0.23 | 1.00 | 34555 | 16310 | |
| Temperature | 0.07 | 0.10 | -0.13 | 0.27 | 1.00 | 34277 | 17137 | |
| Precipitation | -0.03 | 0.06 | -0.16 | 0.10 | 1.00 | 34106 | 16736 | |
| Temperature Seasonality | 0.22 | 0.11 | 0.00 | 0.44 | 1.00 | 21620 | 16683 | |
| Habitat Type: Closed – Open x Migratory Behavior: Absent - Present | 0.06 | 0.15 | -0.23 | 0.36 | 1.00 | 32746 | 17666 | |
| Migratory Behavior: Absent – Present x Territoriality: Non-Territorial - Territorial | **0.81** | **0.27** | **0.29** | **1.33** | 1.00 | 23266 | 16661 | |
| Nest Type: Dome - Primary Cavity x Parental Care: Biparental - Female-Only | 0.00 | 2.51 | -4.93 | 4.92 | 1.00 | 51885 | 13628 | |
| Nest Type: Dome - Secondary Cavity x Parental Care: Biparental - Female-Only | **0.75** | **0.26** | **0.24** | **1.26** | 1.00 | 26215 | 17507 | |
| Nest Type: Dome - Open Cup x Parental Care: Biparental - Female-Only | -0.05 | 0.24 | -0.52 | 0.42 | 1.00 | 25363 | 16961 | |
| Parental Care: Biparental - Female-Only x Clutch Size | 0.02 | 0.25 | -0.47 | 0.50 | 1.00 | 26085 | 17407 | |
| Territoriality: Non-Territorial – Territorial x Social Mating System: Monogamous - Polygynous | -0.42 | 0.29 | -0.99 | 0.14 | 1.00 | 22921 | 17037 | |
| Territoriality: Non-Territorial – Territorial x Wing Length | 0.30 | 0.26 | -0.22 | 0.81 | 1.00 | 17343 | 15567 | |
| Social Mating System: Monogamous – Polygynous x Sexual Size Dimorphism | **0.19** | **0.06** | **0.07** | **0.31** | 1.00 | 31765 | 16631 | |
| Territoriality: Non-Territorial – Territorial x Temperature Seasonality | -0.10 | 0.12 | -0.33 | 0.13 | 1.00 | 21790 | 15856 | |
| Residual Correlation: Female PC2 x Male PC2 | **0.43** | **0.02** | **0.38** | **0.47** | 1.00 | 23392 | 16081 | |

**Legend:** SE is the standard error, L- and U-95% are lower and upper bounds of the 95% credible interval, respectively, $\hat{R}$ is the Gelman-Rubin diagnostic, Bulk and Tail ESS are effective sample sizes, and $R^2$ is the coefficient of determination. The sign × denotes an interaction. Values in bold indicate effects whose credible intervals do not cross zero.

**Table S9.** Contrasts between female and male PC1 in each parameter category and without imputation of missing data in the predictors.

| Parameters | Contrast (Female - Male) | Estimate | l-95% | u-95% |
|---|---|---|---|---|
| Social Pair Bond | Long-Term | 0.08 | -0.03 | 0.19 |
| | Short-Term | **-0.39** | **-0.60** | **-0.19** |
| | Solitary | **-0.56** | **-0.94** | **-0.21** |
| Territoriality × Social Mating System | Non-Territorial : Monogamous | -0.02 | -0.70 | 0.65 |
| | Territorial : Monogamous | -0.01 | -0.11 | 0.09 |
| | Non-Territorial : Polygynous | **-0.55** | **-0.90** | **-0.21** |
| | Territorial : Polygynous | -0.29 | -0.68 | 0.12 |
| Territoriality × Migratory Behavior | Non-Territorial : Migratory | -0.02 | -0.76 | 0.76 |
| | Territorial : Migratory | **-0.47** | **-0.70** | **-0.21** |
| | Non-Territorial : Non-Migratory | **-0.52** | **-0.87** | **-0.15** |
| | Territorial : Non-Migratory | 0.08 | -0.03 | 0.19 |
| Parental Care × Nest Type | Biparental : Primary Cavity | -0.26 | -0.65 | 0.14 |
| | Biparental : Secondary Cavity | **-0.28** | **-0.53** | **-0.05** |
| | Female-Only : Secondary Cavity | -0.10 | -0.75 | 0.51 |
| | Biparental : Dome | -0.16 | -0.37 | 0.06 |
| | Female-Only : Dome | **-0.66** | **-1.23** | **-0.12** |
| | Biparental × Open Cup | **0.19** | **0.05** | **0.33** |
| | Female-Only : Open Cup | **-0.52** | **-0.87** | **-0.17** |
| Migratory Behavior × Habitat Type | Non-Migratory : Closed Habitat | 0.10 | -0.04 | 0.24 |
| | Migratory : Closed Habitat | -0.36 | -0.82 | 0.12 |
| | Non-Migratory : Open Habitat | **-0.21** | **-0.40** | **-0.03** |
| | Migratory : Open Habitat | **-0.46** | **-0.72** | **-0.19** |
| Locomotion Style | Insessorial | -0.17 | -0.40 | 0.06 |
| | Terrestrial | -0.06 | -0.17 | 0.05 |

**Legend:** L- and U-95% are lower and upper bounds of the 95% credible interval, respectively. The sign × denotes an interaction. Values in bold indicate effects whose credible intervals do not cross zero.

**Table S10.** Contrasts between female and male PC2 in each parameter category and without imputation of missing data in the predictors.

| Parameters | Contrast (Female - Male) | Estimate | l-95% | u-95% |
|---|---|---|---|---|
| Social Pair Bond | Long-Term | -0.03 | -0.13 | 0.07 |
| | Short-Term | -0.16 | -0.35 | 0.04 |
| | Solitary | **-0.39** | **-0.73** | **-0.05** |
| Territoriality × Social Mating System | Non-Territorial : Monogamous | -0.01 | -0.64 | 0.65 |
| | Territorial : Monogamous | -0.07 | -0.16 | 0.03 |
| | Non-Territorial : Polygynous | **-0.38** | **-0.69** | **-0.05** |
| | Territorial : Polygynous | 0.10 | -0.28 | 0.49 |
| Territoriality × Migratory Behavior | Non-Territorial : Migratory | -0.02 | -0.74 | 0.70 |
| | Territorial : Migratory | **-0.29** | **-0.52** | **-0.05** |
| | Non-Territorial : Non-Migratory | **-0.35** | **-0.69** | **-0.02** |
| | Territorial : Non-Migratory | 0.00 | -0.11 | 0.10 |
| Parental Care × Nest Type | Biparental : Primary Cavity | -0.12 | -0.50 | 0.24 |
| | Biparental : Secondary Cavity | -0.16 | -0.39 | 0.06 |
| | Female-Only : Secondary Cavity | 0.03 | -0.56 | 0.62 |
| | Biparental : Dome | **-0.24** | **-0.44** | **-0.04** |
| | Female-Only : Dome | -0.24 | -0.76 | 0.29 |
| | Biparental × Open Cup | 0.07 | -0.05 | 0.20 |
| | Female-Only : Open Cup | **-0.34** | **-0.67** | **-0.02** |
| Habitat Type × Migratory Behavior | Non-Migratory : Closed Habitat | 0.03 | -0.10 | 0.16 |
| | Migratory : Closed Habitat | -0.22 | -0.67 | 0.21 |
| | Non-Migratory : Open Habitat | **-0.21** | **-0.38** | **-0.03** |
| | Migratory : Open Habitat | **-0.28** | **-0.53** | **-0.04** |
| Locomotion Style | Insessorial | -0.08 | -0.29 | 0.14 |
| | Terrestrial | -0.09 | -0.19 | 0.01 |

**Legend:** L- and U-95% are lower and upper bounds of the 95% credible interval, respectively. The sign × denotes an interaction. Values in bold indicate effects whose credible intervals do not cross zero.
